# On the utility of Deep Learning for model classification and parameter estimation on complex diversification scenarios

**DOI:** 10.1101/2025.08.27.671290

**Authors:** P.G. Peña, G. Iglesias, E. Talavera, AS. Meseguer, I. Sanmartín

**Author notes:** co-senior authors (AS. Meseguer, I. Sanmartín).

## Abstract

Birth-Death models applied to dated phylogenies are a useful tool to study past diversification dynamics. Parameters in these stochastic models are typically inferred using likelihood-based methods such as Maximum Likelihood Estimation (MLE) or Bayesian Inference. However, these approaches exhibit computational tractability issues in the case of models of moderate to high complexity. One approach to increase model complexity while remaining computationally tractable in the context of birth-death modelling is machine learning. So far, these techniques have been explored in the context of serially-sampled phylogenies (phylodynamics) and trait-dependent birth-death models. Here, we explored the power of Convolutional Neural Networks (CNNs), a type of Deep Learning (DL) method, to solve classification and regression (parameter estimation) tasks under constant-rate and time-homogeneous, rate-variable birth-death models. In particular, we compared six diversification scenarios: Constant Birth-Death, High-Extinction, Mass-Extinction, Diversity-Dependent, Stasis-and-Radiate, and Waxing-and-Waning. We simulated 10, 000 phylogenetic trees under each diversification scenario, which were encoded using a vectorization procedure that captures the topology and branch length information. The encoded trees were used to train or test a set of CNNs models that were designed to tailor three empirical case studies differing in the number of tips. We compared CNNs performance with MLE inference. Our results show that CNNs exhibited classification accuracy levels of 93-78%, whereas maximum likelihood estimation achieved levels of 74-70%. The most difficult scenarios to predict for the CNNs were the high-extinction and mass-extinction scenarios, which were often misidentified as one another. For the regression tasks, mean average errors were comparable between CNNs models and MLE inference, and they also coincided in their difficulty estimating ratio parameters such as mass extinction survival and turnover. Finally, we applied our CNNs to three empirical studies (eucalypts, conifers and cetaceans) and discussed potential shortcomings and future avenues for improvement in the application of deep-learning birth–death modelling approaches.

---

Diversification, defined as the net result of the speciation and extinction processes, is a fundamental process in macroevolution. It plays a critical role in the generation and maintenance of biodiversity, making its study essential for understanding how biodiversity has emerged over geological timescales (Raup et al., 1973; Foote et al., 2007; Morlon et al., 2010; Quental and Marshall, 2010, 2013) and how it has become unevenly distributed across the Earth’s surface or among groups of organisms (Rosenzweig, 1995; Mittelbach et al., 2007; Weir and Schluter, 2007; Rabosky, 2009; Wiens, 2011; Jetz et al., 2012; Rolland et al., 2014; Meseguer and Condamine, 2020).

Despite its inherent importance, the study of diversification has traditionally been a challenge because speciation and extinction are biological processes that typically unfold over millions of years, and extinction, in particular, leaves no trace in the form of extant, living species. Early approaches to understanding diversification relied on inferring diversification rates from the fossil record (Raup et al., 1973; Foote et al., 2007; Ezard et al., 2011). Although promising, this procedure is only suitable for a limited number of organisms, since the fossil record is typically incomplete and sparse for the majority of extant species on Earth, due to gaps and biases in fossil preservation. Dated phylogenetic trees derived from molecular data of present-day species (reconstructed or extant phylogenies) also contain information about evolutionary (speciation, extinction) processes embedded in the distribution of branch lengths and overall topology of the trees (Hey, 1992; Nee et al., 1994b). This finding opened up the possibility of drawing inferences about the diversification process by applying stochastic Birth-Death (BD) models to phylogenies consisting solely of extant taxa.

In its simplest form, BD models include two parameters: the speciation rate (*λ*) that controls the tempo of emergence of new species (birth), and the extinction rate (*µ*) that controls the tempo of elimination of species (death). In addition, a third parameter known as sampling fraction (*ρ*) is often included to account for the fact that not all extant taxa are included in the phylogenetic tree. The simplest BD diversification model is the Constant Birth-Death (CBD), which assumes the rates of *λ* and *µ* are constant over time (Nee et al., 1994a). From this starting point, more complex models with variable rates have been developed. In homogeneous “Time dependent models”, diversification rates can vary over time and these changes affect simultaneously all clades in the phylogeny (Stadler, 2010). Within this type of models, we can differentiate if the time dependence is discrete (Stadler, 2011a; Stadler and Bonhoeffer, 2013; Gavryushkina et al., 2014; Kühnert et al., 2014) or continuous (Rabosky and Lovette, 2008; Morlon et al., 2011b). In “diversity-dependent” models, speciation rates decrease over time as a function of the number of lineages until it reaches a plateau or equilibrium, where speciation equals extinction (Etienne and Haegeman, 2012). Further extensions of BD allow rates to vary as a function of environmental variables (Condamine et al., 2018) or across branches in the phylogeny (Maddison et al., 2007).

All phylogenetic approaches that use BD models to infer past diversification dynamics rely on a common principle: comparing empirical (reconstructed) phylogenies to expectations under various models of diversification (e.g., Stadler, 2011a; Etienne and Haegeman, 2012; Pyron and Wiens, 2013; Höhna et al., 2014; Rolland et al., 2014; Gubry-Rangin et al., 2015; Rabosky et al., 2018; Condamine et al., 2019; Stone and Wolfe, 2021). These approaches differ in the type of representation used to convey the information contained in the phylogenetic tree (edge, critical time, MacPherson et al., 2022), and the statistical framework used to infer the parameters of the model (Morlon, 2014). Within the latter, the most commonly used approaches are Maximum Likelihood and Bayesian Inference. These statistical inferential methods require first to derive the likelihood of the birth-death model and infer the parameter values that maximize the probability of the observed data given the model and set of parameters (Maximum Likelihood, Felsenstein, 1981; Morlon et al., 2011a; Stadler, 2011a), or the posterior probability distribution of the parameters given the data and a prior distribution (Bayesian Inference, Huelsenbeck et al., 2001; Bokma, 2008; Höhna et al., 2019; Vasconcelos et al., 2022). The likelihood of alternative models is compared using different model selection approaches, such as those based on information criteria (Akaike Information Criterion (AIC), Bayesian Information Criterion, weighted AIC (Burnham and Anderson, 2002)), as well as Bayes Factors comparisons, posterior predictive simulations, or cross-validation approaches (Lartillot, 2023).

Despite its wide application, both techniques have important limitations, especially when tackling models of moderate to high complexity. One limitation of likelihood-based methods is that they are often not easily generalizable: for each new model, especially when parameter dependencies change, a new likelihood function must be derived and implemented, often requiring substantial effort. For the most complex models (e.g., SSE-type models), a closed-form likelihood derivation does not exist and numerical integration of Ordinary Differential Equations (ODEs) is necessary; this can introduce problems with numerical approximations and lack of convergence to local optima. Likelihood-based methods can also be affected by computational intractability issues when the diversification model includes a large number of parameters or when the phylogenetic tree includes a large number of taxa (Hinchliff et al., 2015). Another challenge lies in the mechanics of inference itself. Martínez-Gómez et al. (2024) tested five Bayesian methods (Rabosky, 2014; Maliet et al., 2019; Höhna et al., 2019; Barido-Sottani et al., 2020; Maliet and Morlon, 2022; Kopperud and Höhna, 2023) to estimate branch-specific rate changes based on extant-only phylogenetic trees. They concluded that, despite all these methods implementing the same theoretical model (i.e., a rate-heterogeneous BD process), inferences were strongly dependent on the underlying model and assumptions (e.g., anagenetic vs cladogenetic shifts, instantaneous vs. gradual changes). Additionally, Louca and Pennell (2020) demonstrated that, for a given reconstructed phylogenetic tree, there exits an infinity set of “congruent” functions with different values of *λ* and *µ* that yield the same likelihood value. Several solutions have been proposed to address this issue, including reparametrization of the birth-death model (Louca and Pennell, 2020), the use of biologically-informed Bayesian priors (Morlon et al., 2022) and statistical regularization techniques (Andréoletti and Morlon, 2023).

One approach to increase model complexity while remaining computationally tractable in the context of population genetics and phylogenetic diversification is Aproximate Bayesian Computation (ABC, Csilléry et al., 2010; Beaumont, 2019; Bokma, 2010; Janzen et al., 2015). This is a likelihood-free, simulation-based method that searches for parameter values capable of generating data similar to the empirical dataset in terms of summary statistics Summary Statistics (SS), thereby avoiding the need to derive closed-form likelihood functions. However, this procedure has drawbacks, such as the necessity of using a pre-selected set of SS. This can lead to either a significant loss of information if the complexity is overly reduced during the selection process (Robert et al., 2011) or a decrease in accuracy due to high dimensionality when a large number of statistics are used (Blum, 2010; Hartig et al., 2011). Furthermore, it presents an additional limitation because it requires many simulations, which can lead to computational intractability issues.

Machine Learning (ML) based techniques have also begun to be applied as likelihood-free methods for estimating diversification dynamics. ML is a category of artificial intelligence, defined by Arthur Samuel in 1959 as “the field of study that gives computers the ability to learn without being explicitly programmed” (Samuel, 1959). In practice, ML consists in a discipline which encompasses a set of algorithms used to solve classification, regression, clustering, anomaly detection or reinforcement learning problems from a dataset (Alzubi et al., 2018). Depending on the composition of the dataset, there are two fundamental paradigms: supervised machine learning, which involves labelled datasets, or unsupervised machine learning, which deals with unlabelled datasets. Deep Learning (DL) is a subset of machine learning that uses Artificial Neural Networks (ANNs) as parametric learning models. ANNs are able to take raw data as input and extract patterns from it, creating their own high level features, learned from the specific dataset used in during training. This functioning allows them to be broadly flexible about input data structure (Schrider and Kern, 2018) and benefit from high dimensional entry information (LeCun et al., 2015). Convolutional Neural Networks (CNNs) (LeCun et al., 1998) is a well know DL architecture, most commonly applied to the field of pattern recognition within images (Aloysius and Geetha, 2017). The main strengths of CNNs is the ability to capture geometrical features of the data through the convolution mathematical operator; this allows reducing the number of parameters needed to maintain or even improve the model’s performance in comparison with other ML algorithms (e.g., Random Forest (RF) or Feed Forward Neural Network (FFNN)).

ML has been presented as the alternative to likelihood-based methods with the greatest potential in the field of macroevolution, due to their ability to extract high level features directly from raw unstructured data, allowing discrimination of complex diversification scenarios with higher computational efficiency (Bokma, 2006; Sukumaran et al., 2016; Skeels et al., 2023; Lambert et al., 2023; Landis and Thompson, 2025). Bokma (2006) used ML in combination with phylogenetic branching times, summarized in a Principal Component Analysis (PCA), to infer speciation and extinction rates or phenotypic evolution rates. Other authors have used machine learning classification algorithms in combination with phylogenies represented as SS to investigate the effect of ecological traits on organism dispersal rates (Sukumaran et al., 2016), or to explore different eco-evolutionary scenarios (Skeels et al., 2023).

Recently, Voznica et al. (2022), using BD models in a pathogen phylodynamics context, developed a new encoding approach for non-ultrametric phylogenetic trees, known as Compact Bijective Ladderized Vector (CBLV). They combined this tree representation technique with a CNN architecture to perform classification and parameter inference for models of transmission dynamics, proving that this combination has high accuracy and great potential in epidemiology. Lambert et al. (2023) adapted the work of Voznica et al. (2022) to apply DL methods in a macroevolutionary context with trait-dependent diversification models (Maddison et al., 2007), adjusting the CBLV representation to ultrametric trees with associated trait data, the Complete Diversity-reordered Vector (CDV). In this work, parameter regression under a Binary State Speciation and Extinction (BiSSE) model (Maddison et al., 2007) was compared in terms of accuracy with a CBD model (Nee et al., 1994a), using diverse DL architectures (CNN and ANN) and tree representations as inputs (CDV and SS). They concluded that DL inference can reach levels of accuracy in parameter estimation comparable to those obtained with maximum likelihood statistical inference, while being computationally faster by several orders of magnitude. However, one needs also to consider the amount of time required to simulate the datasets and train the model.

Here, we focus on the application of CNNs algorithms for classification (model selection) and regression (parameter estimation) tasks in the context of birth-death models applied to extant phylogenies without associated trait data. We expand CNNs algorithms to discriminate between major diversification scenarios discussed in the literature, some of which have never been addressed in a DL framework, including constant-rate birth-death scenarios, mass extinction scenarios, and time-variable diversification rate scenarios. Our goal is to assess the ability (accuracy and precision) of CNNs to discriminate within a user-defined set of diversification scenarios and to estimate the model parameters under the selected diversification scenario, using the vectorization encoding CDV. We also compared the performance of our CNN approach against Maximum Likelihood Estimation (MLE) inference using whole-tree likelihood methods (Stadler, 2010; Etienne et al., 2012). Finally, we applied CNNs models and MLE to three empirical datasets in order to explore diversification dynamics in real-world phylogenies: a 674-tip phylogeny of eucalypts (Thornhill et al., 2019), a 489-tip phylogeny of conifers (Leslie et al., 2012), and a phylogeny of cetacean mammals with 87 taxa (Steeman et al., 2009). The conifer and cetacean phylogenies have been used as case studies to test the performance of MLE methods for exploring diversification dynamics (Morlon et al., 2011a; Culshaw et al., 2019).

## Materials and Methods

### Macroevolutionary scenarios and Simulations

We explore the potential of CNNs to perform model selection (classification) and estimate the values of diversification parameters (regression) from alternative constant-rate and time-dependent, discrete rate-variable BD scenarios, some of which were already addressed in a previous study using a MLE framework (Sanmartín and Meseguer, 2016). Specifically, we simulated six diversification scenarios to train our CNNs and compare with the MLE method, three with constant rates and three with variable rates: Constant Birth-Death (CBD), High Extinction (HE), Mass Extinction (ME), Diversity Dependent (DD), Stasis and Radiate (SR) and Waxing and Waning (WW). For all these scenarios, we parametrized our simulations with the indirect diversification parameters Relative extinction (a) (Equation (0.1)) and Net Diversification (r) (Equation (0.2)).

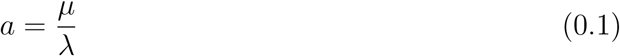

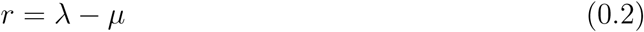

1. Diversification scenarios with constant rates:

a. Constant Birth-Death (CBD): the BD model proposed by Nee et al. (1994b) with constant diversification r and relative extinction a rates, parametrized within the range of values defined by the baseline paleobiological rates.
b. High Extinction (HE): a CBD model but with a high relative extinction rate a, in which at least 80% of speciating lineages go extinct.
c. Mass Extinction (ME): a CBD model interrupted by a Mass Extinction Event (MEE), modelled as a sampling event (i.e. an event that removes a percentage of the standing number of lineages) at a discrete time with magnitude described by parameter *ρ* (survival probability).
2. Diversification scenarios with variable rates:

a. Diversity Dependent (DD): A model in which the speciation rate decreases linearly as a function of the number of lineages, assuming no extinction and with parameter K fixed to 1+ the number of extant taxa.
b. Stasis and Radiate (SR): A discrete time dependent model where diversification remains in stasis until time (t), when it increases sharply.
c. Waxing and Waning (WW): A discrete time dependent model where the diversification rate is initially high (turnover *<* 1) until t, when it decreases dramatically, reaching negative values (turnover *>* 1).

In order to prevent over representation of a particular parameter subspace, we sampled the values for our simulations within prespecified uniform distributions. The bounds of these distributions (Table 1) were selected based on two criteria: (i) ensure that our DL algorithms are “generalist”, that is, the range of simulated values is broad enough to capture multiple diversification trajectories within each scenario, and (ii) use biologically meaningful values. For example, the rate of diversification r was bounded between 0.01 (except for the WW model, see below) and an upper limit of 4.0 diversification events per species per million years; the latter is covering some of the most rapid radiations in animals and plants documented to date (Valente et al., 2010). We also selected the distribution bounds so as to ensure that there was no overlap between different scenarios, e.g., between CBD and HE.

**Table 1.** Uniform distributions (indicated with U) boundaries that were used to sample parameter values for the simulated trees under each of the six diversification scenarios: Constant Birth-Death (CBD), High Extinction (HE), Mass Extinction (ME), Diversity Dependent (DD), Stasis and Radiate (SR) and Waxing and Waning (WW). **r1** and **a1** stand for the values of the diversification rate and turnover parameters, respectively, before the modelled discrete time event **t**. **r0** and **a0** stand for the corresponding values after the discrete time event. *ρ* stands for the proportion of lineages that survives the MEE.

| Model | $r_0$ | $a_0$ | $t$ | $r_1$ | $a_1$ | $\rho$ | $\lambda_0$ |
| --- | --- | --- | --- | --- | --- | --- | --- |
| CBD | U(0.01, 4) | U(0.01, 0.5) | – | – | – | – | – |
| HE | U(0.01, 4) | U(0.8, 0.9) | – | – | – | – | – |
| ME | U(0.01, 1) | U(0.3, 0.8) | U(3, 20) | – | – | U(0.1, 0.3) | – |
| DD | – | – | – | – | – | – | U(0.01, 4) |
| SR | U(0.25, 1.99) | U(0.05, 0.5) | U(3, 20) | U(0.01, 0.1) | U(0.55, 0.95) | – | – |
| WW | U(-0.2, -0.01) | U(1.3, 2) | U(3, 20) | U(0.5, 1.5) | U(0.25, 0.6) | – | – |

To simulate the BD scenarios, we used the R (R Core Team et al., 2013) package TreeSim version 2.4 (Stadler and Stadler, 2019) implementing whole-tree simulations (Stadler, 2011b). In order to create simulated datasets as similar as possible to the empirical cases, simulations were constrained by the number of tips: i.e., each simulation was stopped when it attained the same number of tips as the targeted empirical tree. In the case of models including discrete time events (SR, ME, and WW), we also constrained the crown age of the simulated phylogeny to be at least five million years (M.y.) older than the modelled discrete time event t. The reason for this was to ensure that at least two diversification events separate the crown node and the mass extinction or the rate shift event, assuming a species longevity ranges between 1 and 2.5 M.y.; the average time-to-speciation among plants and animals has been established in 2 Myr (Hedges et al., 2015). We applied a dynamic filter in the simulation script that only annotated the simulated phylogenies if the crown age complies with the criterion mentioned above, thus ensuring that our discrete-time simulated phylogenies do not collapse into a CBD model. This temporal constraint applied to the crown age resulted in a truncated distribution of the simulated parameter value t towards younger ages, but only in the case of SR scenarios (Supplementary Material Figures 15, 16 and 17).

To improve the performance of our DL algorithms (increase accuracy and reduce error), we performed the training phase with phylogenies simulated to a size that matches the number of tips in our empirical case studies. Specifically, for each diversification scenario, we constructed three training datasets of 10, 000 simulated trees with 674 tips (eucalypts, Thornhill et al. (2019)), 489 tips (conifers, Leslie et al. (2012)) and 87 tips (cetaceans, Steeman et al. (2009).

### Phylogenetic tree representation

In order to transform phylogenetic trees into an appropriate structure that can be used as input data by the CNNs, we used the following encoding procedure. First, each tree is rescaled so that the average branch length is 1. Second, each tree is encoded in a one-dimensional vector using the CDV representation.

#### Tree rescaling

Following the same procedure as Voznica et al. (2022) and Lambert et al. (2023), we rescaled the trees to unit average branch length before the encoding, i.e. each branch length on a given tree was divided by the average branch length. We also rescaled the diversification rate under which the tree was simulated by multiplying this value by the tree’s average branch length. We followed the same procedure for the parameter t in the case of the SR, ME, WW models. This was done in order to increase the generality of our DL models and avoid the effect of the time scale, i.e., some simulated trees go deeper in time than others depending on the scenarios and the simulated values. Rescaling ensures that we can apply our trained models on empirical trees with crown ages and underlying diversification rates that differ from those used in the training phase. After the parameter estimation step, we divided the output values by the average branch length, back-scaling them back to the original time scale and ensuring biological interpretability. The only exceptions to this procedure were the relative extinction rate a and *ρ*, as rescaling proportions would have resulted in meaningless parameter values.

#### Encoded tree representation

Phylogenetic trees were encoded using a modified version of the CDV encoding introduced by Lambert et al. (2023) for trait-dependent diversification models, which in turn is a modification on the CBLV procedure developed by (Voznica et al., 2022) for epidemiological phylodynamic models applied to non-ultrametric trees. Our modified CDV procedure involves a preliminary step of internal node reordering, followed by a vectorization step, in which the tree topology is represented as a vector of distances of each node to the root in order to capture the spatial relationship between edges and nodes. Similar to Lambert et al. (2023), our phylogenetic trees are ultrametric, but unlike theirs, there is no tip state data; in fact, our modified CDV (Figure 1), lacks the second vector in the matrix containing the information for the tip states and is almost identical to the one used by Lambert et al. (2023) for the constant-rate birth death model. The main difference with their CDV representation is the removal of the sampling fraction value, and that we skipped the step of completing with zeros in the case of different-size simulated datasets. Specifically, we assumed complete taxon sampling in all simulated trees (sampling fraction =1.0), since our empirical phylogenies were close to 100% taxon sampling, and we did not need to specify the number of tips as encoded information, since each of our simulated phylogenies contained the same number of tips. The remaining steps followed Lambert et al. (2023) and proceeds as follows (Figure 1):

1. **Internal node reordering or ladderization using a diversity criterion**: for each internal node, the sum of the branch lengths of the descending subtree (including the stem branch) is computed, and the internal node with the highest sum is rotated to the left.
2. **Inorder tree transversal and vectorization**: the reordered phylogeny is traversed by recursively starting with the left subtree and for each visited internal node, its distance to the root is added to a vector (Cormen et al., 2009).
3. **Addition of tree height**: we added the value of the tree height as the first vector position.

**Fig. 1.**
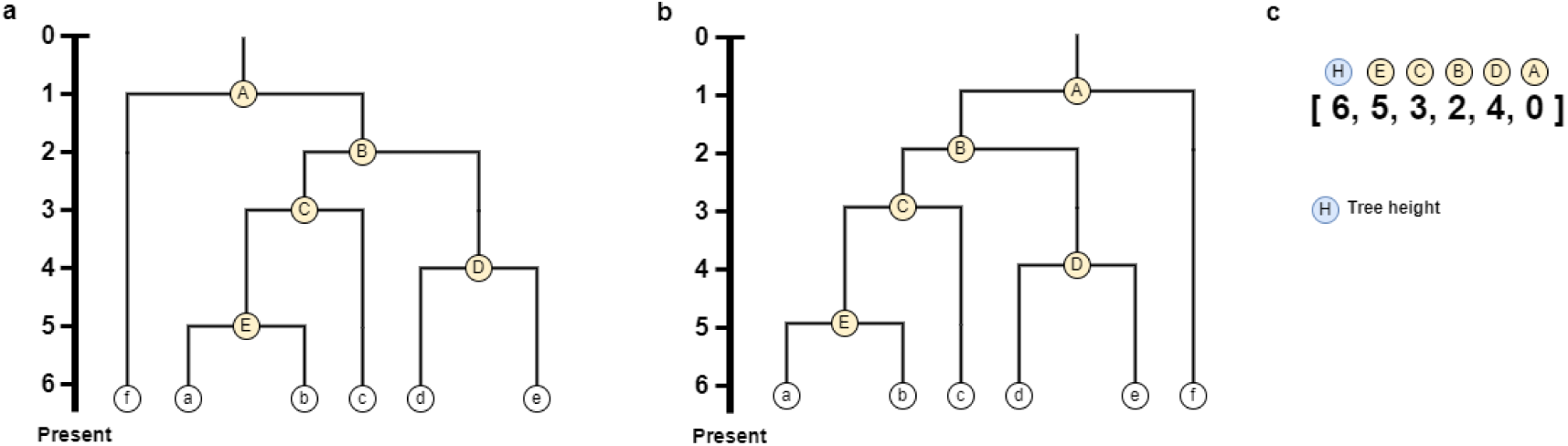
Scheme representing our modified encoding procedure, CDVm with three steps: **a.** Rescaled tree with labelled nodes and tips, and the corresponding edge lengths. **b.** Right-Ladderized tree, in which branches with fewer descendants (or smaller subclades) are placed on the left side of the tree, with larger clades (more descendants) placed on the right. **c.** One-dimensional vector representing the distance of each of the nodes to the root; notice that for ultrametric tips, tip edges are not encoded since they are informed by adding the tree height (H) as the first entry in the encoding vector.

### Model and parameter estimation

Inferring the diversification scenario and rates that are generating one specific phylogenetic tree can be seen as an estimation process. Each phylogenetic tree is defined by a set of parameters that can be either numerical (e.g. a and r) or categorical (i.e., our different diversification scenarios). These parameters can be estimated using different DL models, which will carry out different training procedures depending on the parameters and the scenario. Below, we provide a more detailed explanation of this procedure.

We divided the estimation process in two steps. First, we predict the diversification scenario under which each tree was simulated, using a multi-class classification model: the model ouput is the support for each BD scenario as being the one that generated the tree. Second, the diversification parameters are estimated as a regression problem: we aim to find the combination of values of r, a, t, *ρ* or *λ*_0_ that provides the best fit for the diversification trajectory of a tree under a given scenario. In our parameter estimation approach, we used different DL models for each diversification scenario.

For the DL classification models, we used CDV encoding with a 1-dimensional CNN architecture. CNNs are designed for pattern recognition within images, and are able of extracting features based on local and global patterns. These architectures are particularly appropriate for a type of encoding that maintains local relationships between the vector positions of nodes and edges, such as CDV.

We implemented these approaches in Python 3.8.5, using Tensorflow 1.5.0 (Abadi et al., 2016), Keras 2.2.4 (Chollet et al., 2015) and scikit-learn 0.19.1 (Pedregosa et al., 2011) libraries. For each neural network, we tested several variants differing in the number of layers and neurons, activation function, regularisation process, loss function, and optimiser type; alternative Neural Networks (NNs) were compared in terms of precision and accuracy. For each experiment, the test set was created by randomly shuffling the complete simulation dataset and selecting 20% of it.

#### Diversification scenario estimation using DL models

We designed three models responding to the three empirical examples described in the Introduction. These models differed in their architecture, i.e., the number of convolutional layers increased with the phylogeny’s size (number of tips): CNN for the dataset with trees of 674 tips (CNN-674), CNN for the dataset with trees of 489 tips (CNN-489), and CNN for the dataset with trees of 87 tips (CNN-87) (Figure 2). All three models had one input layer, with the number of input nodes identical to the length of the vector representing the CDV-encoded phylogenetic tree. Unlike Voznica et al. (2022) and Lambert et al. (2023), our architectures lacked the input reshape state, as our trees are ultra-metric and do not include tip state data (i.e. they are encoded as a one-dimensional vector). The input vector flows directly into the convolutional section of the network, which as mentioned above, had different architectures depending of the input size of the data:

- CNN-674: **1)** two 1D convolutional layers of 16 kernels each, with an input size of 3×1, followed by a **2)** max pooling of size 2. Another **3)** two 1D convolutional layers of 32 kernels each with an input size of 3×1 followed by a **4)** max pooling of size 2. **5)** Two additional 1D convolutional layers of 64 kernels each with an input size of 3×1 followed by a **6)** max pooling of size 2 and finally **7)** one 1D convolutional layer with 128 kernels of input size 3×1 and a dropout of 0.3, which helped us to prevent model overfitting.
- CNN-489: The architecture was the same of CNN-674, but we eliminated the steps **5)** and **6)**, as we found that for this smaller phylogeny, a reduced number of convolutional layers achieved equivalent values of precision and accuracy than the more complex CNN-674 architecture.
- CNN-87: The architecture was the same as in CNN-489, except that we removed the steps **3)** and **4)**, given the smaller number of tips.

**Fig. 2.**
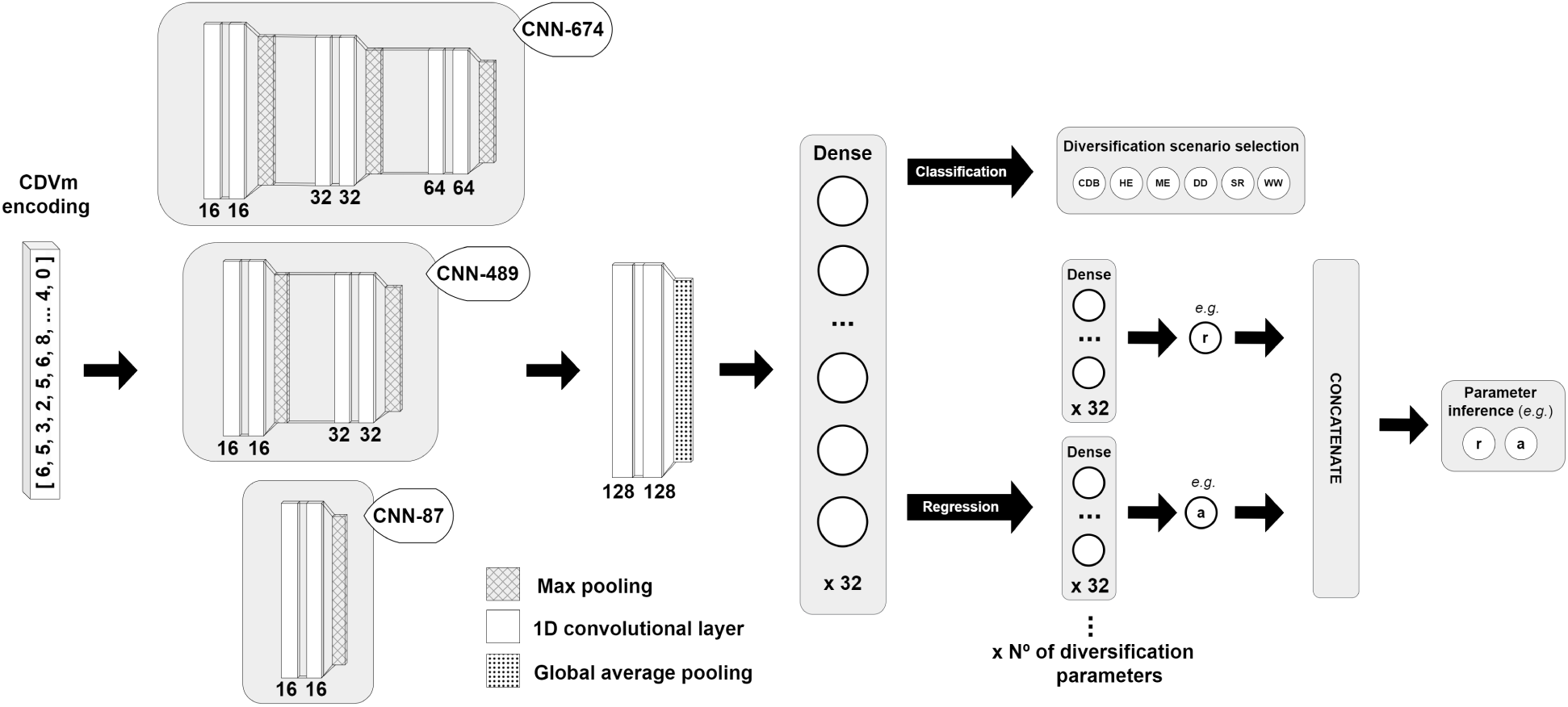
Scheme depicting our different model architectures depending on the task (classification or regression), the number of tips (674, 489 or 87) or the diversification scenario (CBD, HE, ME, DD, SR or WW) they were programmed for. The number at the bottom of each convolutional layer represents the total amount of convolutions and the one under each dense layer represents their total number of neurons.

For the three CNN models, the output of the convolutional part is processed by a a GlobalPoolingAverage1D layer that transforms it to an optimal input for a fully connected ANN with 32 neurons and a dropout of 0.3. All the neurons and kernels of the aforementioned layers used Exponential Linear Unit (ELU) as an activation function.

Finally, we had one output layer with softmax activation function consisting of 6 neurons, one for each diversification scenario. The loss function for these trainings is the *categorical crossentropy*, defined as:

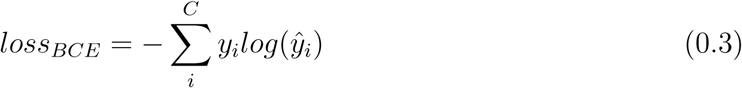

where *C* is the total number of diversification scenarios, *y* is the target scenario and *y*^ is the predicted probability distribution of the scenario.

Of the 10,000 trees simulated for each scenario, we selected ~ 1,000 trees for testing; the 90% remaining trees were used for training, with a 10% of the latter used for validation. All the models were trained with Adam optimizer (Kingma and Ba, 2014) with a learning rate of 0.0001, *β*_1_ of 0.9 and *β*_2_ of 0.999. The batch size used in the training was 128 samples and the models were trained until they did not improve the loss function of the last 50 epochs, using early stopping, with a maximum of 1, 000 epochs.

To explore whether our empirical data-tailored approach performs better than an “all-general” approach, we conducted a subsequent experiment. Following the configuration used for the CNN-674, we tested an alternative approach in which we trained and tested a single classification model on a larger dataset (180,000 simulated trees) that combined all three individual tree-size datasets (674 tips, 489 tips, or 87 tips). To address the problem of different tree sizes, we padded with zero values until reaching the maximum simulated tree size (674 tips).

#### Parameter inference using DL models

To tackle the regression problem, we used CNNs algorithms with phylogenetic trees encoded by CDV. Rather than building a single model to estimate parameters for all scenarios, we created one CNN regression model for each of the six diversification scenarios described above, and for each of the three training datasets (with 674 tips, 489 tips, or 87 tips). While designing 18 regression models might initially appear cumbersome and unnecessary, our preliminary analyses showed that tailoring the training of the CNNs to the specific number of parameters in each model and the size of the simulated trees generated more precise and accurate inferences. Each of these 18 regression models had an architecture similar to the one introduced in the previous section (Figure 2). Unlike the classification model described above, in this architecture the output of the shared 32-neuron dense layer is fed into multiple independent 32-neuron dense layers, each specific to one of the free parameters in the diversification scenario. This allows the model to process each parameter separately with its own dedicated set of weights (Figure 2). Each dense layer was in turn connected to one output layer consisting of a single neuron. The activation function of this neuron for all the parameters was a Leaky Rectified Linear Unit (LeakyReLu), except for the WW scenario, for which we used a linear activation function to allow the corresponding regression model to predict negative values. Finally, the output of each output layer was arranged into a single tensor by performing a concatenation (Figure 2).

The loss function for the parameter inference trainings is the *mean squared error*, defined as:

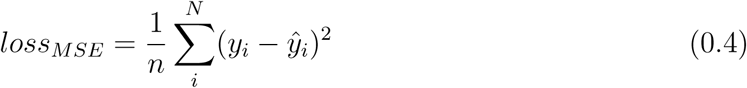

where *N* is the total number of parameters of the model, *y_i_* is the simulated value of the parameter and *y*^*_i_* is the predicted parameter value.

For each diversification scenario, we used the same splitting strategy into training, testing and validation described above for the classification task. All models were trained with Adam optimizer (Kingma and Ba, 2014) with a learning rate of 0.0001, *β*_1_ of 0.9 and *β*_2_ of 0.999. The batch size used in the trainings was 128 samples, and the models were trained as above, stopping when they did not improve the loss function of the last 50 epochs, with a maximum of 1.000 epochs.

We conducted a similar experiment to the one performed for the classification task. Using the configuration of the CNN-674, we trained and tested six regression models (one for each diversification scenario) on a dataset of 30,000 trees that merged all three individual tree-size datasets (10,000 trees each of 674 tips, 489 tips, or 87 tips). Again, to address the issue of different tree sizes, we padded with zero values until reaching the maximum simulated tree size (674 tips).

### Maximum Likelihood Estimation (MLE)

We compared the power and accuracy of our DL models with those obtained using traditional MLE. Specifically, we used whole-tree likelihood inference methods (Stadler, 2011a) implemented in the R package TreePar (Stadler and Stadler, 2015). This package allows modelling constant-rate birth-death scenarios against episodic scenarios with discrete rate shifts in speciation (*λ*), extinction (*µ*) or both, as well as parameter estimation under each of these models. TreePar also allows the user to implement a mass extinction scenario modelled as a constant-rate BD scenario punctuated by a sampling event at discrete time t with magnitude *ρ*, representing the percentage of surviving lineages. Stadler (2011b) demonstrated that it is not possible to distinguish MEEs from rate shifts because both events leave a similar footprint on the phylogeny. Rate shifts are modelled as events that act simultaneously in all lineages, and MEEs are assumed to randomly sample across lineages with magnitude *ρ* (Stadler, 2011a). Additionally, TreePar allows us to infer negative diversification rates, so it can be used for MLE inference under an episodic waxing-and-waning scenario. Unfortunately, TreePar lacks an executable function to evaluate models where diversification rates change as a function of the number of lineages. We used instead the R package DDD v. 5.2.2 (Etienne et al., 2012), implementing model DD 1, which aligns with the diversity-dependent simulation model implemented in TreeSim.

We used TreePar and DDD to perform model selection and parameter estimation under our six diversification scenarios, assuming a sampling fraction of 1.0, in accordance with the simulations. Since MLE is typically more time-consuming than CNN (Voznica et al., 2022; Lambert et al., 2023), we reduced the size of the testing dataset, using 100 simulated trees for each evaluated diversification scenario. To allow a more direct comparison between the two approaches, the same set of 600 simulated trees was employed for additional testing of our DL models. For classification, we used AIC as a statistical measure for model selection. We note, however, that MLE inference in TreePar is not constrained by any prior distribution on parameter values. For some simulated trees, TreePar inferred a “waning-and-waxing” (i.e., decline-then-expansion) scenario in which r is initially negative, a scenario that was not included in our simulation design. To avoid bias against DL, these “not applicable” scenarios were excluded from the evaluation metrics.

### Performance assessment

#### Model selection

Model performance was measured through a set of four evaluation metrics based on the True Positives (TP), True Negaives (TN), False Positives (FP) and False Negatives (FN) of the classification outputs: *accuracy*, *precision*, *recall* and *F1-Score*.

- **Accuracy**: Measures the overall correctness of the model. It is the ratio of the number of correct predictions to the total number of predictions. Its computation is denoted as follows:

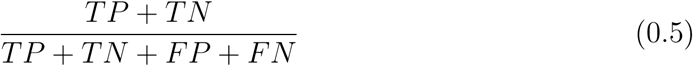

- **Precision** (also known as Positive Predictive Value): measures the correctness of positive predictions. It is the ratio of true positive predictions to the total number of positive predictions. Its computation is denoted as follows:

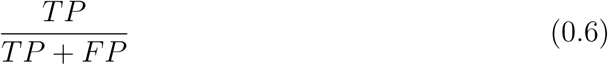

- **Recall** (also known as Sensitivity or True Positive Rate) measures the ability of the model to find all the relevant cases. It is the ratio of true positive predictions to the total number of actual positives. Its computation is denoted as follows:

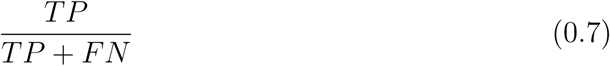

- **F1-Score** is the harmonic mean of precision and recall. It provides a single metric that balances both the concerns of precision and recall, especially useful when you need to find an optimal balance between the two. Its computation is denoted as follows:

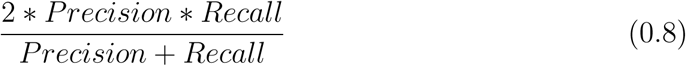

#### Accuracy of parameter inference

To evaluate deviations between the true (“target”) and the predicted parameter values, we employed the Mean Absolute Error (MAE) (Equation (0.9)) and Mean Relative Error (MRE) (Equation (0.10)) measures.

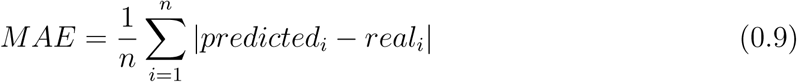

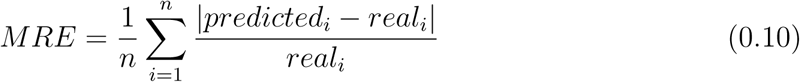

In MLE, the landscape of parameter values explored during inference can be larger than the predefined uniform distributions used for tree simulations. To avoid over-penalizing the MLE approach relative to the CNN models, which were pre-informed with those simulation bounds during the training phase, we followed a similar approach to Lambert et al. (2023). The MLE values inferred outside the simulation ranges were clipped: values below the lower bound were set to the minimum value of our simulation range, and those above the upper bound were set to the maximum simulated value; e.g., if the estimated diversification (r) in the CBD scenario was greater than 4.0, the corresponding value was equated to 4.0. Since DL can potentially predict values outside the simulation ranges (i.e., due to rescaling of trees and parameters), we applied the same strategy to ensure a fair comparison with MLE. Finally, we evaluated the importance of this issue for both MLE and DL by estimating the frequency with which the inferred/predicted values fall outside the simulation ranges.

#### Time efficiency

All experiments (DL and MLE) were conducted on an Intel(R) Xeon(R) Silver 4410Y CPU and an NVIDIA L4 GPU. To assess computational efficiency, we measured the time required for training, testing, and empirical inference for each DL model. For MLE, we measured the time required to perform inference on 100 simulated trees per scenario, as well as the time needed to run the inference on the empirical phylogeny.

### Empirical Illustration

The empirical datasets comprised three different phylogenetic studies. The eucalypts phylogeny is a time-calibrated tree based on the Bayesian phylogeny of Thornhill et al. (2019) (674 tips, excluding outgroups, 83% of sampling fraction), based on two plastid and two nuclear markers. This tree includes taxa from the “true eucalypts” –– genera *Angophora*, *Corymbia*, and *Eucalyptus* –– and was dated using penalized likelihood. Due to very short, near-present branches, we transformed the tree to be binary and ultrametric using the R packages phytools (Revell, 2012) and ape (Paradis et al., 2004). The conifer phylogeny is Leslie et al. (2012) time tree including 489 extant conifer species (we removed the cycads outgroup, c. 80% of sampling fraction), based on two plastid and one nuclear marker, and dated with Bayesian uncorrelated relaxed clocks calibrated with multiple fossil priors. The cetacean time tree is sourced from Steeman et al. (2009) and includes 87 extant species, corresponding to a sampling fraction of 98%. It is based on six mitochondrial and nine nuclear markers; time calibration was performed using several fossils and a penalized likelihood approach.

In order to verify that our empirical data fell within the space covered by our simulations, we performed PCAs with the encoded trees of our empirical examples in their respective simulation datasets.

Importantly, we implemented the following measures to properly account for uncertainty in estimating diversification dynamics from empirical data with our DL classification and regression models:

To mitigate overconfidence in softmax predictions for classification, we applied temperature scaling (Guo et al., 2017). In this approach, a positive scalar parameter *T* (the temperature) is estimated on the validation set by minimizing the negative log-likelihood (NLL) and used to rescale the logits before the softmax transformation. When *T >* 1, the output distribution becomes smoother (increasing entropy and penalizing extreme confidence), while *T <* 1 sharpens it. Importantly, temperature scaling does not change the predicted class, but improves probability calibration.

To quantify model uncertainty, we used Monte Carlo Dropout (MC Dropout) (Gal and Ghahramani, 2016), keeping dropout active at inference and performing multiple forward passes of the empirical tree. This produces a distribution of predictions, from which the mean gives the point estimate and the variance reflects uncertainty. In regression, this distribution also allows calculation of confidence intervals (CIs). MC Dropout provides a practical approximation to Bayesian inference without modifying the network architecture. Importantly, the point estimate remains unchanged, while the spread conveys the model’s confidence.

We combined these measures: for classification, temperature scaling was applied alongside MC Dropout with 10,000 forward passes; for regression, MC Dropout was used with 1,000 forward passes to estimate CIs.

## Results

We assessed the performance of each DL model and MLE for classification (selection among diversification scenarios) and regression (parameter estimation) tasks. After completing the training and evaluation of all models, we inferred the most likely diversification scenario along with the associated parameters for each empirical tree.

For DL, we tested and compared performance among our three sets of models, each corresponding to an empirical case differing in the number of tips (674, 489 and 87), where each set includes one model for classification and six for regression (corresponding to the six diversification scenarios). This allowed us to asses if there are any differences in accuracy and performance related with the phylogenetic tree size. We carried out the same comparison with MLE, using a subsample of our simulation datasets to measure performance and accuracy.

### Simulation time efficiency

On average, simulating the six scenarios required approximately 15 days for the 674-tip dataset, 10 days for the 489-tip dataset, and 16 hours for the 87-tip dataset. However, the WW scenario was consistently the most time-consuming; for example, in the 674-tip dataset, WW accounted for the entire 15-day simulation time, while all other scenarios together were completed in approximately 15 hours. A similar pattern was observed for the other datasets.

### Performance assessment using Deep Learning models

#### Diversification scenario classification

We found that both CNN-674 and CNN-489 report similar overall model accuracy, c. 95%; they also exhibit very similar values for the other metrics. In contrast, for CNN-87, the overall model accuracy drops under 80%. This performance decrease is also reflected across the rest of the metrics, suggesting a relationship between tree size and model performance. Regarding the specific scenarios, the ME case was the most difficult for the model to classify. It was most often confused with HE. Up to a maximum of 21% of ME simulated trees were classified into the wrong scenario in the case of the CNN-674 and CNN-489 models (Supplementary Figure 1). For the remaining scenarios, this percentage dropped to 12% (SR). In the case of the CNN-87, there was a more substantial decrease in precision, with a 53% of misclassified ME trees (Supplementary Figure 1)). Among the other scenarios, this percentage reached a maximum of 35% (HE).

**Table 2.** Classification results: accuracy, precision, recall and F1-Score for each of our DL classification models: CNN-674, CNN-489 and CNN-87.

| DL model | Accuracy | Precision | Recall | F1-Score |
| --- | --- | --- | --- | --- |
| CNN-674 | 0.9398 | 0.9396 | 0.9395 | 0.9393 |
| CNN-489 | 0.9240 | 0.9237 | 0.9237 | 0.9231 |
| CNN-87 | 0.7868 | 0.7832 | 0.7885 | 0.7845 |

The model training time varied between ~ 14 minutes for CNN-674, ~ 24 minutes for CNN-489, and ~ 15 minutes for CNN-87. The test time for the 6,000 test trees ranged between ~ 3 seconds for CNN-674 and less than 1 second for CNN-489 and CNN-87, including the time needed for calibrating the model using temperature scaling.

Merging all tree sizes in a single dataset to train an “all-general” model resulted in accuracy and precision values lower than those obtained with the data-tailored models in the case of the CNN-674 and CNN-489, but higher for the CNN-87 (Supplementary Material Table 7). The model training time was ~ 1 hour.

#### Diversification parameter estimation

Our regression results (Table 3 and Supplementary Table 1) show that the errors in parameter estimation (MAE) are generally similar across all parameters, except for t. This parameter exhibits values at least two orders of magnitude higher than the other parameters, which is expected given that it has the largest numerical values and that MAE is an absolute error metric. Regarding the DL regression models, we found a similar behaviour than for the classification models: CNN-674 and CNN-489 reported similar MAE values, whereas CNN-87 is associated with higher values. Supplementary Table 1 shows the corresponding MRE values, which show similar patterns to the MAE metric.

**Table 3.** MAE for DL and MLE. The error measures for DL are obtained after testing with 1,000 phylogenetic trees for each of the 18 DL regression models (one for each of the six different diversification scenarios and each of our three empirical datasets: 674, 489, and 87 tips). For MLE, the error measures are based on the inference results of 100 trees for each diversification scenario and tip size.

| Tree size | Estimation type | Parameter |  |  |  |  |  |  |
| --- | --- | --- | --- | --- | --- | --- | --- | --- |
| | | r0 | a0 | t | r1 | a1 | $\rho$ | $\lambda_0$ |
|  |  | Constant Birth-Death |  |  |  |  |  |  |
| 674 tips | DL | 0.1274 | 0.0722 | - | - | - | - | - |
|  | MLE | 0.1166 | 0.0688 | - | - | - | - | - |
| 489 tips | DL | 0.1266 | 0.0736 | - | - | - | - | - |
|  | MLE | 0.1510 | 0.0805 | - | - | - | - | - |
| 87 tips | DL | 0.2291 | 0.1080 | - | - | - | - | - |
|  | MLE | 0.3459 | 0.1433 | - | - | - | - | - |
|  |  | High Extinction |  |  |  |  |  |  |
| 674 tips | DL | 0.2462 | 0.0191 | - | - | - | - | - |
|  | MLE | 0.1875 | 0.0188 | - | - | - | - | - |
| 489 tips | DL | 0.2678 | 0.0260 | - | - | - | - | - |
|  | MLE | 0.2019 | 0.0185 | - | - | - | - | - |
| 87 tips | DL | 0.6111 | 0.0315 | - | - | - | - | - |
|  | MLE | 0.3088 | 0.0246 | - | - | - | - | - |
|  |  | Mass Extinction |  |  |  |  |  |  |
| 674 tips | DL | 0.0391 | 0.0673 | 4.8021 | - | - | 0.0496 | - |
|  | MLE | 0.0338 | 0.0574 | 0.9089 | - | - | 0.0496 | - |
| 489 tips | DL | 0.0384 | 0.0776 | 4.3575 | - | - | 0.0495 | - |
|  | MLE | 0.0264 | 0.0580 | 0.7131 | - | - | 0.0543 | - |
| 87 tips | DL | 0.0368 | 0.0939 | 5.2705 | - | - | 0.0483 | - |
|  | MLE | 0.0567 | 0.1230 | 1.8258 | - | - | 0.0757 | - |
|  |  | Diversity Dependent |  |  |  |  |  |  |
| 674 tips | DL | - | - | - | - | - | - | 0.0622 |
|  | MLE | - | - | - | - | - | - | 0.0631 |
| 489 tips | DL | - | - | - | - | - | - | 0.0664 |
|  | MLE | - | - | - | - | - | - | 0.0700 |
| 87 tips | DL | - | - | - | - | - | - | 0.1640 |
|  | MLE | - | - | - | - | - | - | 0.1800 |
|  |  | Stasis and Radiate |  |  |  |  |  |  |
| 674 tips | DL | 0.0700 | 0.1132 | 3.5182 | 0.0217 | 0.0961 | - | - |
|  | MLE | 0.0579 | 0.0899 | 1.2234 | 0.0375 | 0.1693 | - | - |
| 489 tips | DL | 0.0573 | 0.1011 | 3.2836 | 0.0573 | 0.1011 | - | - |
|  | MLE | 0.0468 | 0.0848 | 1.5694 | 0.0347 | 0.1644 | - | - |
| 87 tips | DL | 0.0709 | 0.1076 | 2.9354 | 0.0204 | 0.0917 | - | - |
|  | MLE | 0.1439 | 0.1869 | 2.0699 | 0.0415 | 0.1964 | - | - |
|  |  | Waxing and Waning |  |  |  |  |  |  |
| 674 tips | DL | 0.0361 | 0.1471 | 2.7317 | 0.1054 | 0.0826 | - | - |
|  | MLE | 0.0351 | 0.2117 | 0.3356 | 0.1407 | 0.1621 | - | - |
| 489 tips | DL | 0.0304 | 0.1349 | 2.7102 | 0.0999 | 0.0863 | - | - |
|  | MLE | 0.0386 | 0.1796 | 0.3904 | 0.0974 | 0.1550 | - | - |
| 87 tips | DL | 0.0343 | 0.1340 | 3.0987 | 0.1278 | 0.0892 | - | - |
|  | MLE | 0.0425 | 0.1815 | 0.6091 | 0.1451 | 0.1678 | - | - |

The percentage of predicted parameter values, across all scenarios and parameter values, that fell outside the simulated ranges was below 7% for the 674-tip dataset, *<* 9% for the 489-tip dataset, and *<* 24% for the 87-tip dataset (Table 6). We notice that for parameters *ρ* in the ME scenario, a0 and a1 in SR and a0 and a1 in the WW scenario, the CNN models predicted the mean value regardless of the target value (Supplementary Figures 9, 10 and 11).

The mean training time among our three empirical-specific set of regression CNNs models ranged between 1.5 minutes ±14 seconds for the models trained with 674-tip phylogenies, 2.2 minutes ±30 seconds for the models trained 489-tip phylogenies, and 1.9 minutes ±55 seconds for the models trained with 87-tip phylogenies. For performing the testing phase, the mean time varied between 0.77 ± 1.11 seconds for the models trained with 674-tip phylogenies, 0.32 ± 0.07 seconds for the models trained 489-tip phylogenies, and 0.26 ± 0.06 seconds for the models trained with 87-tip phylogenies.

Regarding the “all-general” models, results followed a similar pattern to the classification task. MAE values were generally higher than those obtained with the data-tailored models in the case of the regression models trained with 674 tips and 489 tips, but lower for the models trained with 87 tips (Supplementary Material Table 8). On average, the time spent training each model was approximately 20 minutes.

#### Performance assessment using Maximum Likelihood Estimation Diversification scenario classification

For the classification problem, MLE accuracy was between 71-74% for the three empirically-based simulation datasets (674 tips, 489 tips and 87 tips) under the set of tested scenarios; i.e, excluding the “not applicable” cases (Table 4). Similar values were reported for the other metrics, irrespective of tree size. Regardless the specific scenario and tree-size dataset, the percentage of missclasified scenarios in MLE was generally higher than in the DL models (Supplementary Figures 2 and 1). The false negative rates (“type II error”), based on the AIC criterion, was higher than in the CNNs models. The highest rate of false positives (“type I error”) was found in the ME scenarios, followed by SR. Interestingly, the highest rate of erroneous assignment to other, “not applicable” scenarios corresponded also to SR. (Supplementary Table 2).

**Table 4.** Classification results: accuracy, precision, recall and F1-Score for MLE.

| Number of tips | Accuracy | Precision | Recall | F1-Score |
| --- | --- | --- | --- | --- |
| 674 | 0.7313 | 0.8089 | 0.7330 | 0.7327 |
| 489 | 0.7406 | 0.8063 | 0.7418 | 0.7394 |
| 87 | 0.7153 | 0.7267 | 0.6941 | 0.6948 |

#### Diversification parameter estimation

Table 3 lists the MAE obtained after testing TreePar with 100 phylogenetic trees for each diversification scenario and each empirically-based simulation dataset. Overall, the MAE values with MLE were similar to those obtained with theCNN models for the simulated phylogenies with 674 and 489 tips, whereas this error was slightly higher for the simulated phylogenies with 87 tips. The only exception was the MAE for the t parameter, which was considerably lower for MLE inference than the corresponding values for the CNN regression models (table 3). As in the CNN models, we observe a pattern of increasing MAE values with decreasing tree size. Supplementary Table Table 1 shows the corresponding MRE values, which show similar patterns.

The percentage of predicted parameter values, across all scenarios and parameter values, that fell outside the simulated ranges reached values as high as 75% for the 674-tip datasets, 75% (489-tip datasets), and 93% for the 87-tip datasets (Table 5). Considering the specific parameters, those with the highest percentage of inference outside the bounds were *ρ* in the ME scenario, a0 and a1 in SR, and r0, a0 and a1 in the WW scenario.

**Table 5.** DL and MLE predicted diversification parameters for the empirical data: for eucalypts, conifers, and cetaceans. 95% confidence intervals estimated with MC dropout are also included for the DL estimates.

|  |  | r0 | a0 | t | r1 | a1 |
| --- | --- | --- | --- | --- | --- | --- |
| Eucalypts | DL | 0.3062<br>[0.225–0.4125] | 0.2479<br>[0.200–0.34] | 1.4708<br>[0.45–2.50] | 0.0085<br>[0.01–0.02] | 0.6855<br>[0.62–0.72] |
|  | MLE | -0.2610 | 1.2180 | 5.0141 | 0.0688 | 0.9620 |
| Conifers | DL | 0.0610<br>[0.05–0.11] | 0.3093<br>[0.200–0.375] | 0.4939<br>[0.2–0.8] | 0.0051<br>[0.004–0.008] | 0.7167<br>[0.64–0.74] |
|  | MLE | 0.0338 | 0.7631 | 24.5506 | 0.0125 | 0.7031 |
| Cetaceans | DL | 0.1728<br>[0.11–0.24] | 0.2265<br>[0.16–0.28] | 1.8094<br>[1–2.75] | 0.0163<br>[0.01–0.022] | 0.7380<br>[0.7–0.78] |
|  | MLE | 0.0620 | 0.0919 | 2.7882 | 0.0802 | 0.5202 |

MLE inference time ranged from ~ 12 hours for the 674-tips dataset, to ~ 10 hours for the 489 tips dataset and ~ 5 hours for the 87 tips dataset.

### Empirical Illustration

Principal Component Analysis (PCAs), performed over the vectorization (CDV) of the three training datasets and the empirical phylogenies, showed that the empirical data was well nested within the simulation space. Together, First Axis of Principal Component Analysis (PCA1) and Second Axis of Principal Component Analysis (PCA2), explained 98% of the data variance for eucalytps, 98% for conifers, and 91% for cetaceans (Supplementary Figures 3, 4 and 5).

Our trained classification DL models predicted –– using 10,000 forward passes with MC Dropout on each empirical tree –– that the diversification scenario for eucalypts, conifers, and cetaceans was Stasis and Radiate (SR), with the mean of the scaled-softmax probabilities equal to 1.0, 0.97, and 0.59, respectively. For cetaceans, the CBD scenario also showed relatively high scaled-softmax probabilities, with a mean of 0.48. Regression results, including the mean of the predicted diversification parameters and associated 95% confidence intervals estimated from 1,000 MC Dropout forward passes, are reported in Table 5.

The MLE classified the eucalypt phylogeny under a WW scenario, with an AIC of 2762.5. The next best-supported model was the ME scenario, with an AIC of 2771.7, resulting in a ΔAIC of less than 10. In the case of conifers, the empirical phylogeny was classified as a SR scenario with an AIC of 3573.5; the next selected scenario, ME, has an AIC= 3575.0. Finally, the cetacean phylogeny was classified as a SR scenario with AIC = 556.5, compared to CBD with AIC= 557.6. The parameter estimation results are shown in Table 5

Inference time of each empirical case varied from ~ 13 minutes using MLE and ~ 1 minute using DL (including the time spent for performing MC Dropout in classification and regression) for the eucalypt phylogenetic tree, to ~ 40 minutes using MLE and ~ 35 seconds with DL for the conifer tree, and ~ 1 minute for MLE and ~ 34 seconds with DL for the cetacean phylogeny.

## Discussion

In this study, we present an approach based on Deep Learning, CNNs, for classification and regression tasks in a birth-death modelling framework. Within the macroevolutionary field, Lambert et al. (2023) already explored the potential of DLs for parameter estimation applied to constant-rate (CBD) and clade(trait)-dependent (BiSSE) diversification scenarios. Lambert’s study was centred on the regression task, i.e., performing parameter value estimations. In order to develop a widely applicable, generalist model, these authors simulated millions of phylogenetic trees, with different number of tips. Parameter estimation was based on simulations sampled from uniform distributions with relatively narrow ranges, for example, parameter *λ* in the CBD was bounded by a uniform distribution between 0.01 and 0.5. Here, we propose a different approach that is empirically-driven. We tailored our simulation design to a specific tree size informed by the empirical case. This allowed us to explore a wider range of diversification scenarios, i.e., six time-constant and time-variable models, using broader parameter prior distributions (e.g., r was bounded between 0.01 and 4.0). In doing so, we aimed to capture the stochasticity of biological processes observed in empirical data, while keeping it within biologically realistic thresholds (Valente et al., 2010).

A data-tailored DL model might seem less efficient than the more generalist approach employed by Lambert et al. (2023) and Voznica et al. (2022). However, it presents several advantages. By training our models with data that closely mirrors the structure of the empirical tree, we significantly reduced the number of simulations required to maximise the accuracy of our CNNs, i.e., we used 10, 000 simulations per scenario compared to four million in Lambert et al. (2023); initial trial tests demonstrated that increasing the number of simulations by one order of magnitude (100, 000 simulations per scenario) did not increase the CNNs performance and accuracy relative to the 10, 000 simulated datasets. We also managed to decrease the time spent in model training. Simulation time (15 hours for the largest dataset) was lower than in Lambert et al. (2023) (~ 2 days) across all scenarios with the exception of Waxing and Waning (WW) (~ 14 days simulating for the largest dataset). We consider that the inclusion of this rarely explored birth–death scenario, even though computationally demanding, is a strength of our lightweight simulation approach (i.e., it would be computationally intractable to simulate one million trees).

The “all-general” approach, which combines all three individual tree-size datasets to train the DL models, did not achieve the performance levels of the data-tailored models for the 674-tip and 489-tip cases, despite being trained on a larger set of simulated trees. Moreover, the average training time for the classification task was considerably shorter for the data-tailored models (17 minutes versus one hour), with a similar pattern observed for the regression models (1 minute versus 22 minutes). We therefore argue that tailoring the DL model to the problem is a more efficient strategy, yielding comparable levels of performance with substantially shorter training times and fewer simulations (e.g., 60,000 compared with 180,000 trees for the classification task).

Our results showed that DLs trained on vector-encoded phylogenies produced accuracy levels for classification tasks that exceeded 93% for the largest simulated phylogenies, and 78% for the 87-tip phylogenies. These results were replicated when testing the same validation dataset used for MLE (Supplementary Table 3). Furthermore, our DL models outperformed MLE for model selection: TreePar and DDD reached maximum levels of accuracy and precision that were lower than 74% and 80%, respectively, regardless of the simulated tree size (71% and 72% for the 87-tip phylogenies).

In terms of discriminating among diversification scenarios, our DL models precision results indicate that HE and ME were the most difficult scenarios to predict, and that they were often misidentified as one another. This confusion stems, in part, from the fact that we have used a “relaxed” concept of mass extinction. In the paleontological literature, it is typically assumed that the percentage of lineages removed during the MEE is at least 90% of the standing diversity at that point in time (Sepkoski, 1982; Culshaw et al., 2019). In contrast, following other studies tackling ME in plants (Antonelli and Sanmartín, 2011; May et al., 2016), we allowed this percentage to be lower, by implementing a uniform distribution for the *ρ* parameter (survival probability) bounded as U(0.3, 0.1). This definition allowed us to detect less “strict” MEEs, in which up to 30% lineages could survive the MEE. This percentage corresponds to extinction levels estimated for the Terminal Eocene Event at the Late Eocene–Early Oligocene boundary, when a dramatic cooling of climates extirpated evergreen plant lineages that once formed part of the Holarctic boreotropical flora (Zachos et al., 2001; Antonelli and SANMARTin, 2011; Culshaw et al., 2019). However, when our simulations adopt values in the lower range of the uniform distribution of the *ρ* parameter(around 0.3), the simulated phylogenetic trees resemble those simulated under the HE scenario, which would explain the CNN confusion.

In the case of MLE, there is very little confusion between HE and ME. However, ME and SR scenarios were the most frequently selected scenarios across all tree sizes and irrespective of the simulated scenario, i.e., high Type I and type II errors (Supplementary fig. 2). Laurent et al. (2014) also found that TreePar tended to overestimate the frequency of MEE. This percentage of type II error for the SR scenario is higher for the 87-tip dataset, in agreement with other studies (Stadler, 2011a; Sanmartín and Meseguer, 2016) reporting reduced statistical power for MLE methods applied to small trees. Furthermore, between 20% and 30% of our simulated SR trees were inferred by TreePar as corresponding to a “waning-and-waxing” scenario, i.e., a decline-then-expansion scenario in which r is initially negative. This scenario was not included in our simulation set. This may appear to give CNNs an advantage over MLE when comparing overall accuracy and precision. However, when correcting for this bias – by excluding these scenarios – we still find a lower accuracy in the classification of scenarios when compared with CNN models. These observations suggest that when there is prior knowledge to constrain the set of plausible diversification scenarios, DL approaches might be more advantageous than MLE. Further studies comparing other MLE diversification methods are needed to establish the generality of this conclusion.

Regarding the regression task, comparing the MAE and MRE values from our DL regression models with those from MLE revealed that our CNNs achieved comparable levels of performance for all parameters in the simulated scenarios, regardless of the number of tips in the phylogenies (Table 3 and Supplementary Table 1). We observed these same levels of performance when our CNNs were tested with the same validation datasets used for MLE (Supplementary Table 4). The only exception was parameter t in the discrete-change diversification scenarios ME, SR, and WW, which was associated with higher MAE and MRE values in the DL regression models compared to MLE (Table 3; Supplementary Tables 4 and 1).

The distribution of the error metric values (*predicted_i_* − *real_i_*) around each parameter was well centred. The mean and median values were very close to zero, and there were often symmetrical levels of underestimation and overestimation with respect to the simulated values (Supplementary Figure 6, Figure 7 and Figure 8). Nevertheless, when we further explored the correlation between the error and the target (simulated) values (Supplementary Material figures 9, 10 and 11), we observed a tendency of the DL models to return the mean values for certain parameters: the survival probability *ρ* in the ME scenario and the turnover a in the discrete-time scenarios SR and WW. We notice that parameters of this type, with a ratio form, were also the most often inferred outside the simulation bounds by MLE (Supplementary Material Table 5). For example, parameter a1 (turnover in the distant past) in the SR scenario is inferred outside the bounds in ~ 93% of cases for the 87-tip dataset, ~ 63% for the 489-tip dataset, and ~ 70% for the 674-tip dataset. This skewed inference produces unusual error distributions, often forming an “X” pattern around the mean. This occurs because values inferred above the upper bound are truncated (“clipped”) at the upper bound; as target values approach the upper bound, the apparent error decreases (Supplementary Material Figures 12, 13 and 14). These results suggest that parameters challenging DL estimation are also problematic for likelihood-based inference. Indeed, compound (indirect) parameters such as turnover are often reparametrized (e.g., equating it with extinction) because they introduce additional dependencies between *λ* and *µ* (Stadler and Bonhoeffer, 2013), and induce unusual prior distributions than complicate MCMC convergence (Höhna, 2014). Given that both DL and MLE methods encounter difficulties in estimating parameters of this type (turnover and *ρ*), we believe that it is preferable to predict the mean under a biologically informed prior rather than to provide a point estimate outside this prior, which could be biologically unrealistic.

The lower model performance of CNN models with small phylogenies (*<* 100 tips), as evidenced by the 87-tip dataset, is in line with the results of (Lambert et al., 2023), who demonstrated that the error in parameter estimation using CDV-informed CNNs, increased as the number of tips in the phylogeny decreased. The limited phylogenetic information captured by small trees when encoded as whole-tree vectors, relative to larger trees, could be alleviated by using alternative encodings such as Summary Statistics (SS). Several studies have reported good precision values using this encoding combined with ANNs, even in the case of small trees (Voznica et al., 2022; Lambert et al., 2023; Lajaaiti et al., 2023). Since SS is a representation with no spatial or temporal hierarchical order, we believe that combining simple ML algorithms, such as random forest or decision trees, with SS encoding would be a more efficient approach to tackle the problem of small-size trees in the context of diversification studies. Another solution for DL inference applied to small trees could be the one adopted in PhyloCNN (Perez and Gascuel, 2024), which uses node-feature encoding and might be less dependent on tree size.

In the case of the empirical phylogenies, the DL and MLE approaches gave different results regarding the classification and/or regression tasks. We emphasize that the goal of our empirical analyses is to illustrate how the method and trained networks can be used on empirical data, since there is no “ground truth” for “real-world” phylogenies. The only possible comparison is therefore against other sources of empirical evidence. The eucalypt phylogeny was classified as a SR scenario by DL with an acceleration of diversification rates starting around 1.5 M.y. In contrast, MLE selected the WW scenario with a rate shift towards declining rates around 5 M.y. The SR scenario with a rapid expansion in the last 1.5 million years captured by the CNN is in agreement with a recent systematic study (Thornhill et al., 2019). These authors inferred that several species-rich sections within subgenus *Symphyomyrtus* (563 species), belonging to genus *Eucalyptus*, originated in the Pleistocene, in the last million years. Many of these species have been described based on morphological characters, as “morphospecies”, and are subtended in Thornhill et al. (2019)’s tree by near zero branch lengths. Our CNNs were seemingly able to capture information from these very short distal branches, identifying “polytomies” as (nearly simultaneous) diversification events that occurred very close to the present. The manner in which phylogenetic information is encoded and processed using CNNs-CDV might explain the difference with MLE, which infers the timing of the rate shift nearly four million years earlier. MLE represents the tree as a vector of branching times, each of them representing a bifurcation event. The CNNs-CDV models introduced here also encode this information, but additionally, through the ladderization procedure, they recorded for each node in the tree which daughter lineage gave rise to more descendants (Figure 1). In other words, the CDV encoding might be able to capture the degree of balancedness in the empirical tree, in this case caused by a very recent radiation, whereas MLEvectorization of branching times does not seem to.

A similar explanation might apply to the conifer phylogeny. Both DLs and MLE classified this tree as a SR scenario, but the time of the rate shift was estimated quite differently: c. 25 M.y. by MLE and 0.5 M.y. by DL. Likelihood inference performed with Bayesian inference methods (Culshaw et al., 2019; May et al., 2016) identified the Middle Oligocene as a period of sudden increase in diversification rates, associated with global cooling, which could be mistaken for a mass extinction event (Culshaw et al., 2019). This is also in agreement with the fossil record (Leslie et al., 2012). Instead of this event, our DL model captures the diversification events occurring in the last 0.5 million years, evidenced by several clades within genera *Juniperus*, *Hesperocyparis*, *Abies*, *Picea*, and specially *Pinus*, which are subtended by near-zero branch lengths in the original phylogeny (Leslie et al., 2012). It is interesting to note that many species of *Eucalyptus* and *Pinus* are fire-adapted –– some even require fire to germinate –– and that fire-prone habitats became widespread in the Late Pleistocene-Holocene, which might explain the rapid radiation events during this period (Thornhill et al., 2019).

For the cetacean tree, we do not have a plausible explanation for the CBD versus the SR model identified as the best model by the CNN and MLE inference, respectively. Nevertheless, the difference from the next best model, CBD, is not substantial for either DL –– where the mean scaled-softmax probabilities were ~60% for SR and ~40% for CBD –– or for MLE, where the AIC of CBD is only 1.0 unit larger than that of the SR model. It might be that none of these scenarios adequately represents the diversification dynamics of this group. Morlon et al. (2011a) detected a pattern of lineage-specific diversification, with some cetacean clades experiencing diversity decline (whales) and other clades undergoing lineage expansion towards the present (dolphins). The tree-wide diversification scenarios examined in this study, in which all lineages undergo simultaneous expansion or decline, will not be able to capture this clade-dependent waxing-and-waning-and-waxing dynamics.

Regarding empirical inference, an important advantage of the DL approach introduced here is computational efficiency. In terms of computational time, classification and parameter estimation was one order of magnitude longer with MLE than with our DL models for the eucalypt dataset.

Previous studies have shown that when the training data does not accurately represent the real data, learning-based methods can behave “poorly”, as the trained model has never encountered data similar to the empirical (Clemmensen and Kjærsgaard, 2022). In previous works (Voznica et al., 2022; Lambert et al., 2023), some “sanity checks” are typically performed in order to verify that the empirical phylogenetic trees are similar to the simulated ones, such as using SS metrics to compare empirical and simulated values, or employ PCA methods. We used here the PCA approach, which confirmed that the vectorized empirical phylogenies fall within the eigen vector space defined by the simulated trees. However, studies using other data types (e.g., simulated DNA and protein alignments) under both DL (Trost et al., 2024) and MLE (Guindon et al., 2010; Höhler et al., 2022) approaches have found significant, yet poorly understood, differences between simulated and empirical (“real”) data. We believe that this issue also extends to phylodynamics and the simulation of birth-death trees. We constrained our simulated trees to match the number of tips in the empirical tree, attempting to create simulated datasets as similar as possible to the empirical cases. However, we believe that this approach may be not sufficient to increase our confidence in the predictions.

It has been long known that real-world phylogenies tend to be more unbalanced than trees generated by birth-death models (Kirkpatrick and Slatkin, 1993). This imbalance can result from several factors which are not accounted for by the simple CBD models used to simulate phylogenetic data, such as incomplete lineage sorting, lineage-dependent rate shifts, or “clade”-specific mass extinction events (Blum and Fraņcois, 2006; Sanmartín and Meseguer, 2016; Fischer et al., 2023). We tested the “unbalance” hypothesis by calculating the Colless Index (Bartoszek et al., 2021) for each of the empirical and simulated phylogenies, using the R package *treebalance* (Fischer et al., 2023). Supplementary Material Figure 18, Figure 19 and Figure 20 show that the empirical value falls outside the 95% distribution of values from the simulated datasets and that this result is consistent across all empirical phylogenies. The tree-balance analysis suggests that the set of tree-wide BD models analyzed here is not sufficient to capture the complexity of diversification patterns represented by the empirical trees. One factor likely to affect the comparison between empirical and simulated trees is the presence of different diversification scenarios –– i.e., independent diversification trajectories for different clades –– in the original phylogenies. This may be the case for the conifer and eucalypt trees, where the original studies, using evidence from the fossil record, point to different selective pressures per clade, driven by varying environmental conditions associated with their geographical distributions (Leslie et al., 2012; Thornhill et al., 2019). Similarly, Morlon et al. (2011a) proposed that independent waxing-and-waning histories for different cetaceans clades could explain their diversification trajectory, based on MLE inference and the fossil record.

An interesting solution to the issue of the lack of similarity between empirical and simulated phylogenies is to use unsupervised NNs, informed by fossil data (Hauffe et al., 2024). However, the application of this approach might be limited to those few groups of extant organisms with an abundant fossil record. Another solution is to increase the set of simulated scenarios by including more complex models, for example, multiple-rate shift episodic models, clade-dependent diversification, fossilized birth death processes, or trait evolutionary models, including discrete or continuous adaptive models. This increasing complexity, however, runs the risk of introducing computational tractability issues. One future research avenue that could enable us to tackle different evolutionary processes and scenarios with CNNs approaches is the development of more sophisticated phylogenetic representation/encoding languages, as well as more powerful DL algorithms to process them. A tentative first step was taken by Lajaaiti et al. (2023), who combined a graph-based representation of phylogenetic trees with Graphical Neural Networks (GNN). This combination, however, resulted in a poor performance due to “over-smoothing” and “hop neighbourhood” problems. We think these issues could be solved by adopting more powerful DL architectures. For example, a recent study has shown good results for parameter estimation for some macroevolutionary models using an ensemble of different deep-learning architectures and graph-based tree encoding (Qin et al., 2024). Nevertheless, it remains to be explored whether this approach could be extended to other birth-death models.

## Conclusions

Deep Learning, specifically Convolutional Neural Network approaches, have become increasingly popular in evolutionary biology, especially within the fields of demographic inference, species delimitation problems, and epidemiology. In contrast, to date, there are only few applications of CNNs in the macroevolutionary field. In this study, we explore the performance of CNNs for classification and parameter estimation in the context of time-homogeneous, constant-rate or rate-variable birth-death models, including mass extinction events and discrete-time rate shifts. Using broad uniform distributions to capture biological stochasticity in diversification parameters, we demonstrated that DL is able to perform significantly better than MLE for all classification tasks, and exhibits a better performance for most regression tasks. Unlike previous efforts, we added model selection as a previous step before parameter estimation, ensuring that the CNN model architecture used for the subsequent regression task matches the selected diversification scenario. Ours is the first exploration of DL to diversification dynamics involving time-variable birth-death scenarios. Future research avenues should include establishing robust procedures to compare simulated and real phylogenetic trees and to develop more efficient encoding languages that can capture the complexity of real-world diversification dynamics.

## Acknowledgements

This study was funded through projects PID2020-120145GA-I00 to ASM and PID2019-108109GB-I00 to IS by the MCIN/AEI/10.13039/501100011033/ and EURFD ‘A way of making Europe’, and project PID2023-153023NB-I00 to ASM and IS by the MCIN/AEI/10.13039/501100011033/ and European Union NextGenerationEU/PRTR. The grants “*Ayuda para la contratación de ayudante de investigación*” (PEJ-2021-AI/AMB-21780) funded by the Fondo Social Europeo and the *Programa Operativo de Empleo Juvenil y la Iniciativa de Empleo Juvenil* of the Comunidad de Madrid to ASM and PGP, and by a doctoral grant of the Comunidad de Madrid (PIPF-2022/ECO-25483) to PGP, supervised by ASM and IS. We thank Pablo Ariño for evaluating the code and providing valuable feedback that greatly improved this work. We thank Manolo Fernandez Perez for his valuable feedback and insightful comments on this manuscript. We thank the research group KNOwledge Discovery and Information Systems (KNODIS) for the compute infrastructure.

## Data Availability

All the original data and scripts necessary to reproduce the analyses reported in this study can be accessed through the Dryad Digital Repository: Dryad Repository.

**Sharelink**: This is the Sharelink

**GitHub**: deep-birth-death

## Author contributions

Pablo G. Peña: conceptualization, investigation, formal analysis, writing (original draft); Guillermo Iglesias: investigation, formal analysis, writing (original draft); Edgar Talavera: resources; Isabel Sanmartín: conceptualization, investigation, formal analysis, writing (original draft), supervision; Andrea Sánchez Meseguer: Funding acquisition, conceptualization, writing (review and editing), supervision.

## Supplementary Material

**Table 1.**
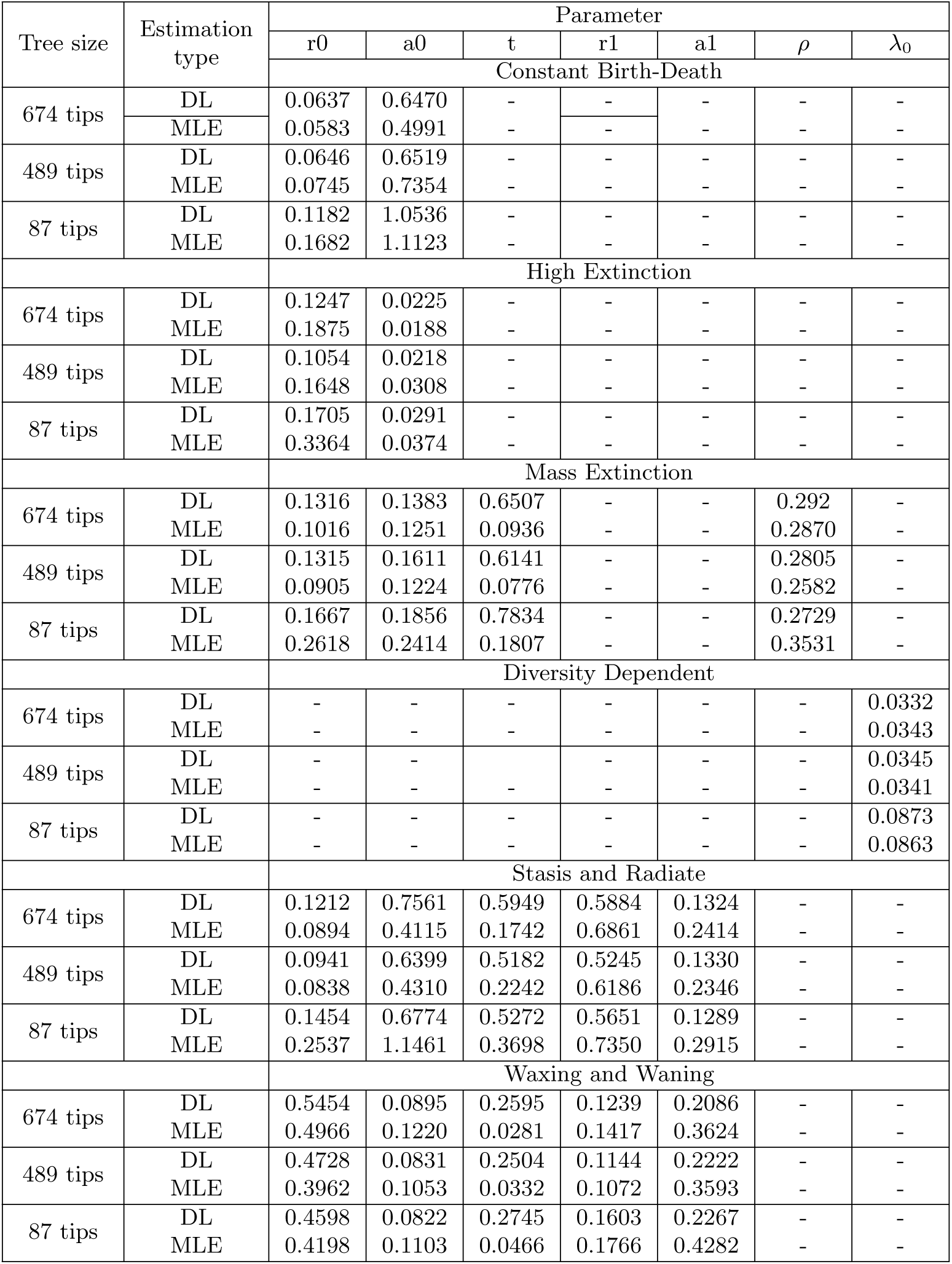
MRE for DL and MLE. The error measures for DL are obtained after testing with 1,000 phylogenetic trees each of the the 18 DL regression models (one for each of the 6 different diversification scenarios and for each of our three empirical datasets: 674, 489, and 87 tips). For MLE the error measures are based on the inference results of 100 trees for each diversification scenario and tip size

| Tree size | Estimation type | Parameter |  |  |  |  |  |  |
| --- | --- | --- | --- | --- | --- | --- | --- | --- |
| | | r0 | a0 | t | r1 | a1 | $\rho$ | $\lambda_0$ |
|  |  | Constant Birth-Death |  |  |  |  |  |  |
| 674 tips | DL | 0.0637 | 0.6470 | - | - | - | - | - |
|  | MLE | 0.0583 | 0.4991 | - | - | - | - | - |
| 489 tips | DL | 0.0646 | 0.6519 | - | - | - | - | - |
|  | MLE | 0.0745 | 0.7354 | - | - | - | - | - |
| 87 tips | DL | 0.1182 | 1.0536 | - | - | - | - | - |
|  | MLE | 0.1682 | 1.1123 | - | - | - | - | - |
| High Extinction |  |  |  |  |  |  |  |  |
| 674 tips | DL | 0.1247 | 0.0225 | - | - | - | - | - |
|  | MLE | 0.1875 | 0.0188 | - | - | - | - | - |
| 489 tips | DL | 0.1054 | 0.0218 | - | - | - | - | - |
|  | MLE | 0.1648 | 0.0308 | - | - | - | - | - |
| 87 tips | DL | 0.1705 | 0.0291 | - | - | - | - | - |
|  | MLE | 0.3364 | 0.0374 | - | - | - | - | - |
| Mass Extinction |  |  |  |  |  |  |  |  |
| 674 tips | DL | 0.1316 | 0.1383 | 0.6507 | - | - | 0.292 | - |
|  | MLE | 0.1016 | 0.1251 | 0.0936 | - | - | 0.2870 | - |
| 489 tips | DL | 0.1315 | 0.1611 | 0.6141 | - | - | 0.2805 | - |
|  | MLE | 0.0905 | 0.1224 | 0.0776 | - | - | 0.2582 | - |
| 87 tips | DL | 0.1667 | 0.1856 | 0.7834 | - | - | 0.2729 | - |
|  | MLE | 0.2618 | 0.2414 | 0.1807 | - | - | 0.3531 | - |
| Diversity Dependent |  |  |  |  |  |  |  |  |
| 674 tips | DL | - | - | - | - | - | - | 0.0332 |
|  | MLE | - | - | - | - | - | - | 0.0343 |
| 489 tips | DL | - | - | - | - | - | - | 0.0345 |
|  | MLE | - | - | - | - | - | - | 0.0341 |
| 87 tips | DL | - | - | - | - | - | - | 0.0873 |
|  | MLE | - | - | - | - | - | - | 0.0863 |
| Stasis and Radiate |  |  |  |  |  |  |  |  |
| 674 tips | DL | 0.1212 | 0.7561 | 0.5949 | 0.5884 | 0.1324 | - | - |
|  | MLE | 0.0894 | 0.4115 | 0.1742 | 0.6861 | 0.2414 | - | - |
| 489 tips | DL | 0.0941 | 0.6399 | 0.5182 | 0.5245 | 0.1330 | - | - |
|  | MLE | 0.0838 | 0.4310 | 0.2242 | 0.6186 | 0.2346 | - | - |
| 87 tips | DL | 0.1454 | 0.6774 | 0.5272 | 0.5651 | 0.1289 | - | - |
|  | MLE | 0.2537 | 1.1461 | 0.3698 | 0.7350 | 0.2915 | - | - |
| Waxing and Waning |  |  |  |  |  |  |  |  |
| 674 tips | DL | 0.5454 | 0.0895 | 0.2595 | 0.1239 | 0.2086 | - | - |
|  | MLE | 0.4966 | 0.1220 | 0.0281 | 0.1417 | 0.3624 | - | - |
| 489 tips | DL | 0.4728 | 0.0831 | 0.2504 | 0.1144 | 0.2222 | - | - |
|  | MLE | 0.3962 | 0.1053 | 0.0332 | 0.1072 | 0.3593 | - | - |
| 87 tips | DL | 0.4598 | 0.0822 | 0.2745 | 0.1603 | 0.2267 | - | - |
|  | MLE | 0.4198 | 0.1103 | 0.0466 | 0.1766 | 0.4282 | - | - |

**Fig. 1.**
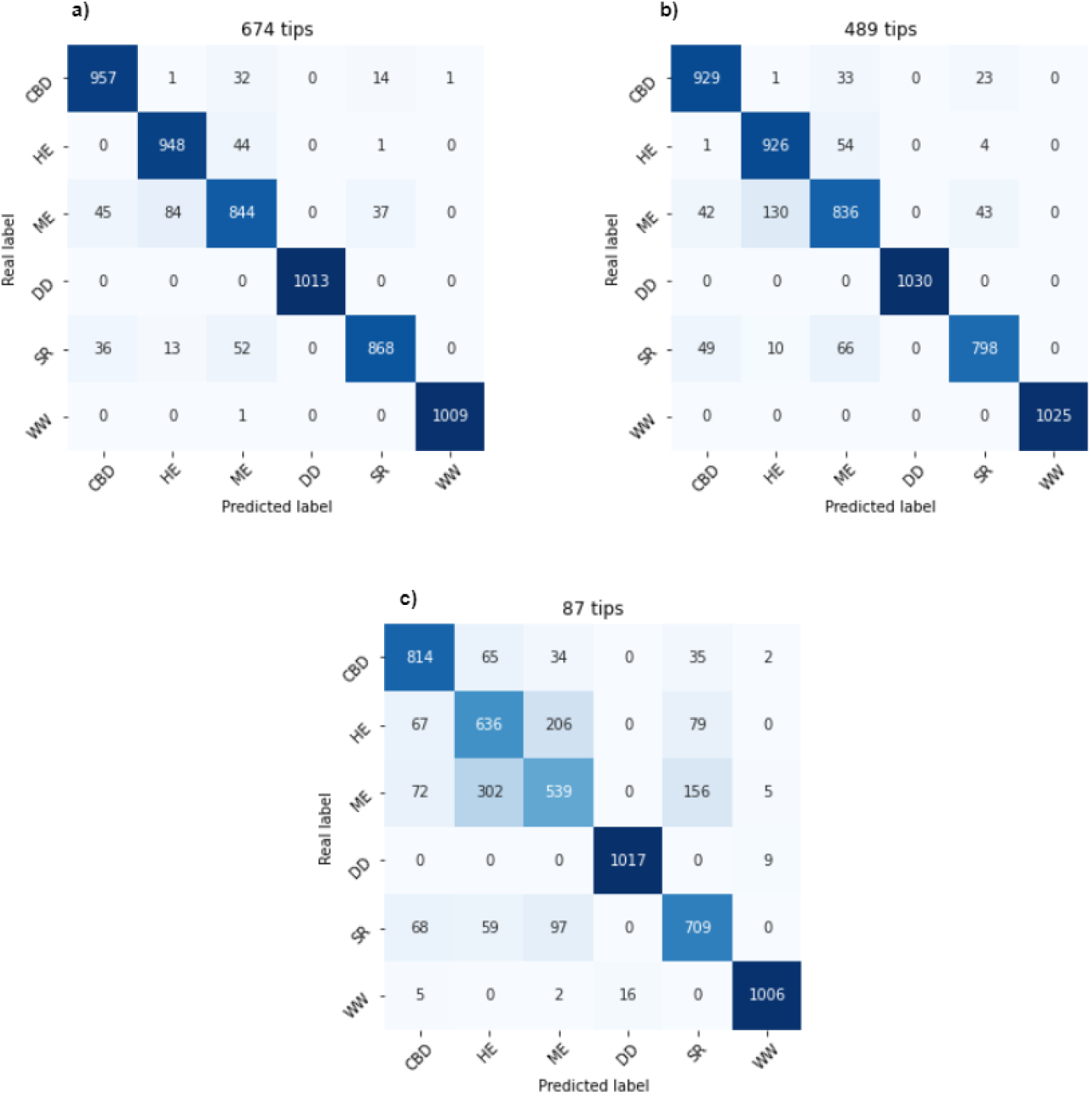
Confusion matrices generated by testing the DL classification models with ~ 1,000 simulated phylogenetic trees of each diversification scenario. Rows indicate the real labels (simulated diversification scenario) and columns, the predicted labels. Each cell value represents the number of predictions falling into the corresponding real–predicted label combination, with higher diagonal values indicating better classification performance. (a) Confusion matrix for the model trained on simulated phylogenetic trees with 674 tips. (b) Confusion matrix for the model trained on simulated trees with 489 tips. (c) Confusion matrix for the model trained on simulated trees with 87 tips.

**Fig. 2.**
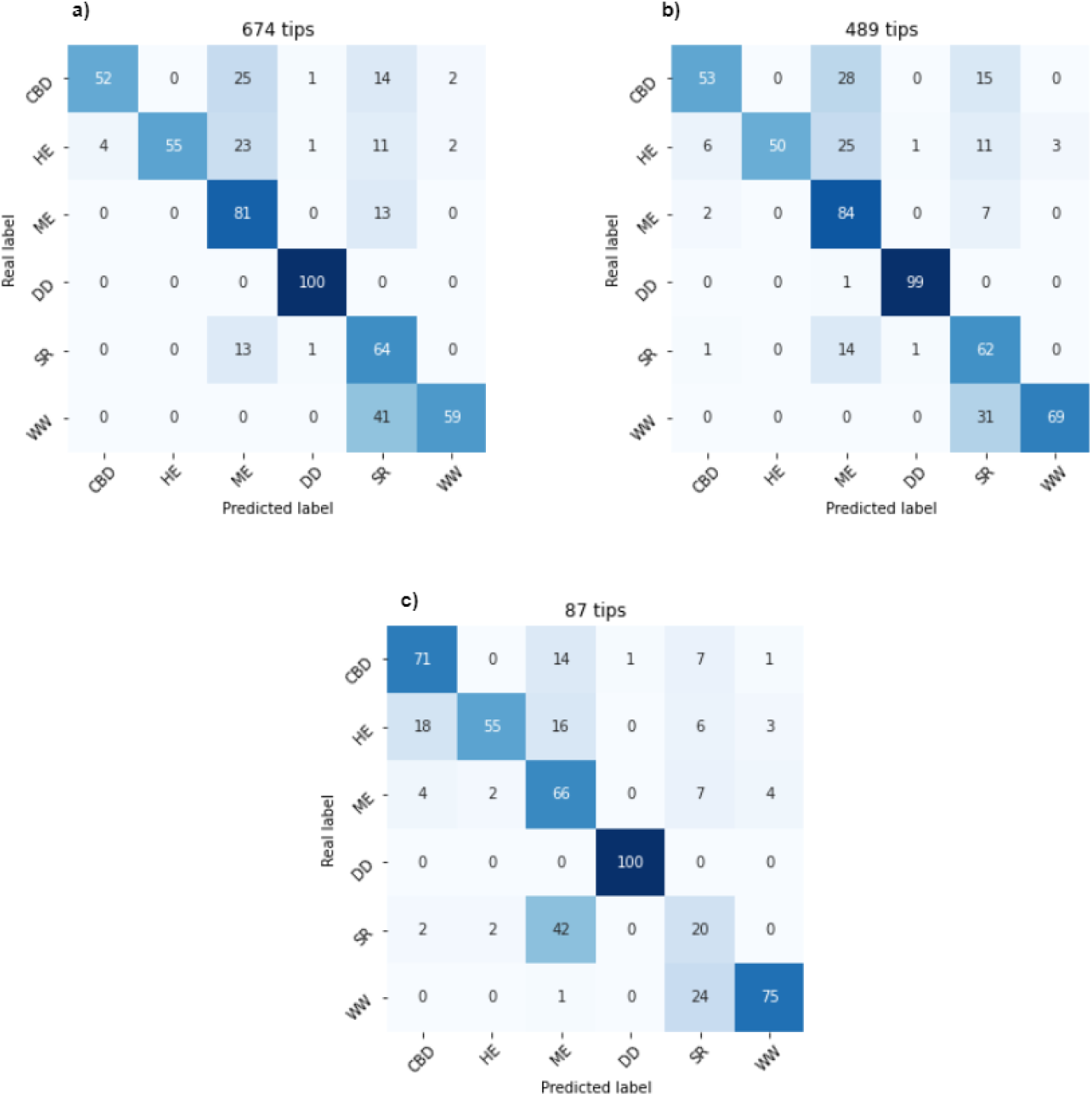
Confusion matrices generated by performing MLE on 100 simulated phylogenetic trees for each diversification scenario, using AIC as the model selection criterion. All other conventions follow Fig. 1. (a) Confusion matrix for simulated phylogenetic trees with 674 tips. (b) Confusion matrix for simulated trees with 489 tips. (c) Confusion matrix for simulated trees with 87 tips. For some scenarios, the row values do not sum 100 because some TreePar occasionally predicted scenarios outside our predefined set; see Supplementary Table 2.

**Table 2.** Number of “not applicable” cases reported by TreePar for each diversification scenario and number-of-tips combination. “Not applicable” refer to cases in which the selected scenario is not within the predefined simulation set.

|  | 674 tips | 489 tips | 87 tips |
| --- | --- | --- | --- |
| CBD | 6 | 4 | 6 |
| HE | 4 | 2 | 2 |
| ME | 6 | 7 | 17 |
| DD | 0 | 0 | 0 |
| SR | 22 | 22 | 34 |
| WW | 0 | 0 | 0 |

**Table 3.** Classification results showing accuracy, precision, recall, and F1-score for each of our DL classification models (CNN-674, CNN-489, and CNN-87), evaluated on the same simulation dataset that was used for inference with MLE.

| Number of tips | Accuracy | Precision | Recall | F1-Score |
| --- | --- | --- | --- | --- |
| 674 | 0.9250 | 0.9243 | 0.9250 | 0.9236 |
| 489 | 0.9350 | 0.9348 | 0.9350 | 0.9342 |
| 87 | 0.8133 | 0.8133 | 0.8133 | 0.8125 |

**Table 4.**
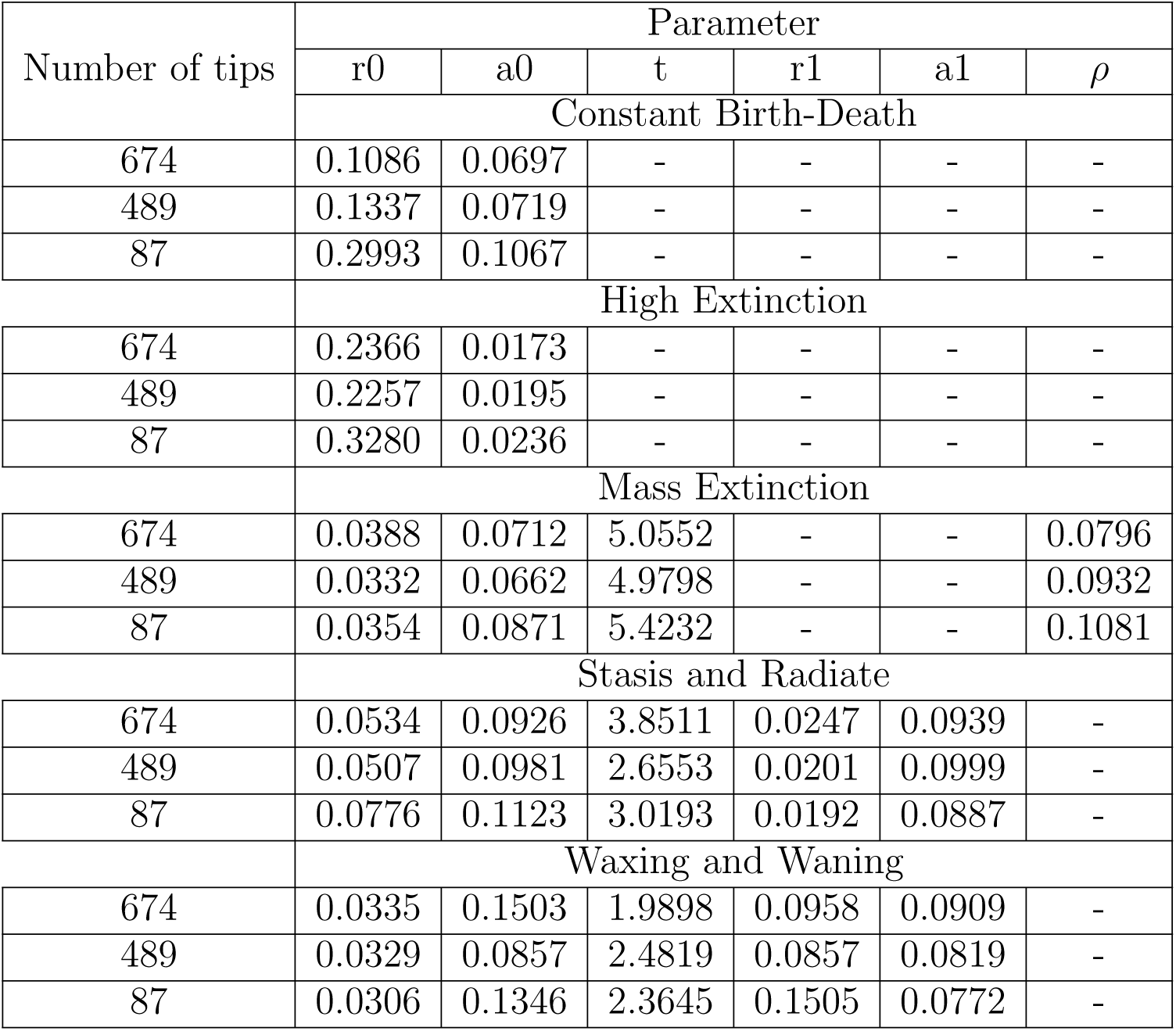
Parameter estimation results: Mean Absolute Error (MAE) obtained by each of our DL regression models (CNN-674, CNN-489, and CNN-87), evaluated on the same simulation dataset that was also used for inference with MLE.

**Table 5.** Percentage of inferred parameters that fall outside the simulation range in MLE. “674 t.” refers to the simulated phylogenies of 674 tips, “489 t.” to those of 489 tips, and “87 t.” to the simulated phylogenies with 87 tips. “Clipped” indicates the total percentage of inferred parameters whose value is outside the simulation range.

| 674 t. | r0 | a0 | t | r1 | a1 | $\rho$ | $\lambda_0$ | 489 t. | r0 | a0 | t | r1 | a1 | $\rho$ | $\lambda_0$ |
| --- | --- | --- | --- | --- | --- | --- | --- | --- | --- | --- | --- | --- | --- | --- | --- |
|  | Constant Birth-Death |  |  |  |  |  |  |  | Constant Birth-Death |  |  |  |  |  |  |
| Clipped | 1% | 17% | - | - | - | - | - | Clipped | 2% | 16% | - | - | - | - | - |
| High Extinction |  |  |  |  |  |  |  | High Extinction |  |  |  |  |  |  |  |
| Clipped | 5% | 21% | - | - | - | - | - | Clipped | 8% | 24% | - | - | - | - | - |
| Mass Extinction |  |  |  |  |  |  |  | Mass Extinction |  |  |  |  |  |  |  |
| Clipped | 0% | 16% | 5 | - | - | 30% | - | Clipped | 0% | 11% | 1 | - | - | 40% | - |
| Diversity Dependent |  |  |  |  |  |  |  | Diversity Dependent |  |  |  |  |  |  |  |
| Clipped | - | - | - | - | - | - | 3% | Clipped | - | - | - | - | - | - | 3% |
| Stasis and Radiate |  |  |  |  |  |  |  | Stasis and Radiate |  |  |  |  |  |  |  |
| Clipped | 3% | 23% | 8% | 38% | 68% | - | - | Clipped | 7% | 24% | 8% | 33% | 63% | - | - |
| Waxing and Waning |  |  |  |  |  |  |  | Waxing and Waning |  |  |  |  |  |  |  |
| Clipped | 58% | 50% | 1% | 10% | 75% | - | - | Clipped | 46% | 41% | 2% | 6% | 75% | - | - |

| **87 t.** | r0 | a0 | t | r1 | a1 | $\rho$ | $\lambda_0$ |
| --- | --- | --- | --- | --- | --- | --- | --- |
|  | Constant Birth-Death |  |  |  |  |  |  |
| Clipped | 7% | 38% | - | - | - | - | - |
| High Extinction |  |  |  |  |  |  |  |
| Clipped | 10% | 55% | - | - | - | - | - |
| Mass Extinction |  |  |  |  |  |  |  |
| Clipped | 0% | 31% | 10 | - | - | 54% | - |
| Diversity Dependent |  |  |  |  |  |  |  |
| Clipped | - | - | - | - | - | - | 4% |
| Stasis and Radiate |  |  |  |  |  |  |  |
| Clipped | 32% | 70% | 28% | 68% | 93% | - | - |
| Waxing and Waning |  |  |  |  |  |  |  |
| Clipped | 43% | 43% | 3% | 23% | 79% | - | - |

**Table 6.** Percentage of inferred parameters that fall outside the simulation range in DL. “674 t.” refers to the simulated phylogenies of 674 tips, “489 t.” to those of 489 tips, and “87 t.” to the simulated phylogenies with 87 tips. “Clipped” indicates the total percentage of inferred parameters whose value is outside the simulation range.

| **674 t.** | r0 | a0 | t | r1 | a1 | $\rho$ | $\lambda_0$ |
| --- | --- | --- | --- | --- | --- | --- | --- |
|  | Constant Birth-Death |  |  |  |  |  |  |
| Clipped | 4% | 1% | - | - | - | - | - |
| High Extinction |  |  |  |  |  |  |  |
| Clipped | 6% | 1% | - | - | - | - | - |
| Mass Extinction |  |  |  |  |  |  |  |
| Clipped | 0% | 0.1% | 8% | - | - | 0% | - |
| Diversity Dependent |  |  |  |  |  |  |  |
| Clipped | - | - | - | - | - | - | 1.5% |
| Stasis and Radiate |  |  |  |  |  |  |  |
| Clipped | 3% | 0% | 8% | 0% | 0% | - | - |
| Waxing and Waning |  |  |  |  |  |  |  |
| Clipped | 0.7% | 0% | 6% | 1.4% | 0% | - | - |

| **489 t.** | r0 | a0 | t | r1 | a1 | $\rho$ | $\lambda_0$ |
| --- | --- | --- | --- | --- | --- | --- | --- |
|  | Constant Birth-Death |  |  |  |  |  |  |
| Clipped | 3.5% | 0% | - | - | - | - | - |
| High Extinction |  |  |  |  |  |  |  |
| Clipped | 4% | 0.4% | - | - | - | - | - |
| Mass Extinction |  |  |  |  |  |  |  |
| Clipped | 0.1% | 0% | 7% | - | - | 0% | - |
| Diversity Dependent |  |  |  |  |  |  |  |
| Clipped | - | - | - | - | - | - | 2% |
| Stasis and Radiate |  |  |  |  |  |  |  |
| Clipped | 4.2% | 0% | 8.8% | 0.1% | 0% | - | - |
| Waxing and Waning |  |  |  |  |  |  |  |
| Clipped | 1% | 0% | 8.9% | 1.1% | 0% | - | - |

| **87 t.** | r0 | a0 | t | r1 | a1 | $\rho$ | $\lambda_0$ |
| --- | --- | --- | --- | --- | --- | --- | --- |
|  | Constant Birth-Death |  |  |  |  |  |  |
| Clipped | 5.7% | 0% | - | - | - | - | - |
| High Extinction |  |  |  |  |  |  |  |
| Clipped | 9.7% | 0% | - | - | - | - | - |
| Mass Extinction |  |  |  |  |  |  |  |
| Clipped | 0% | 0% | 8.5% | - | - | 0% | - |
| Diversity Dependent |  |  |  |  |  |  |  |
| Clipped | - | - | - | - | - | - | 3.1% |
| Stasis and Radiate |  |  |  |  |  |  |  |
| Clipped | 5% | 0% | 12% | 0.5% | 0% | - | - |
| Waxing and Waning |  |  |  |  |  |  |  |
| Clipped | 1.5% | 0.7% | 4.4% | 23% | 0% | - | - |

**Fig. 3.**
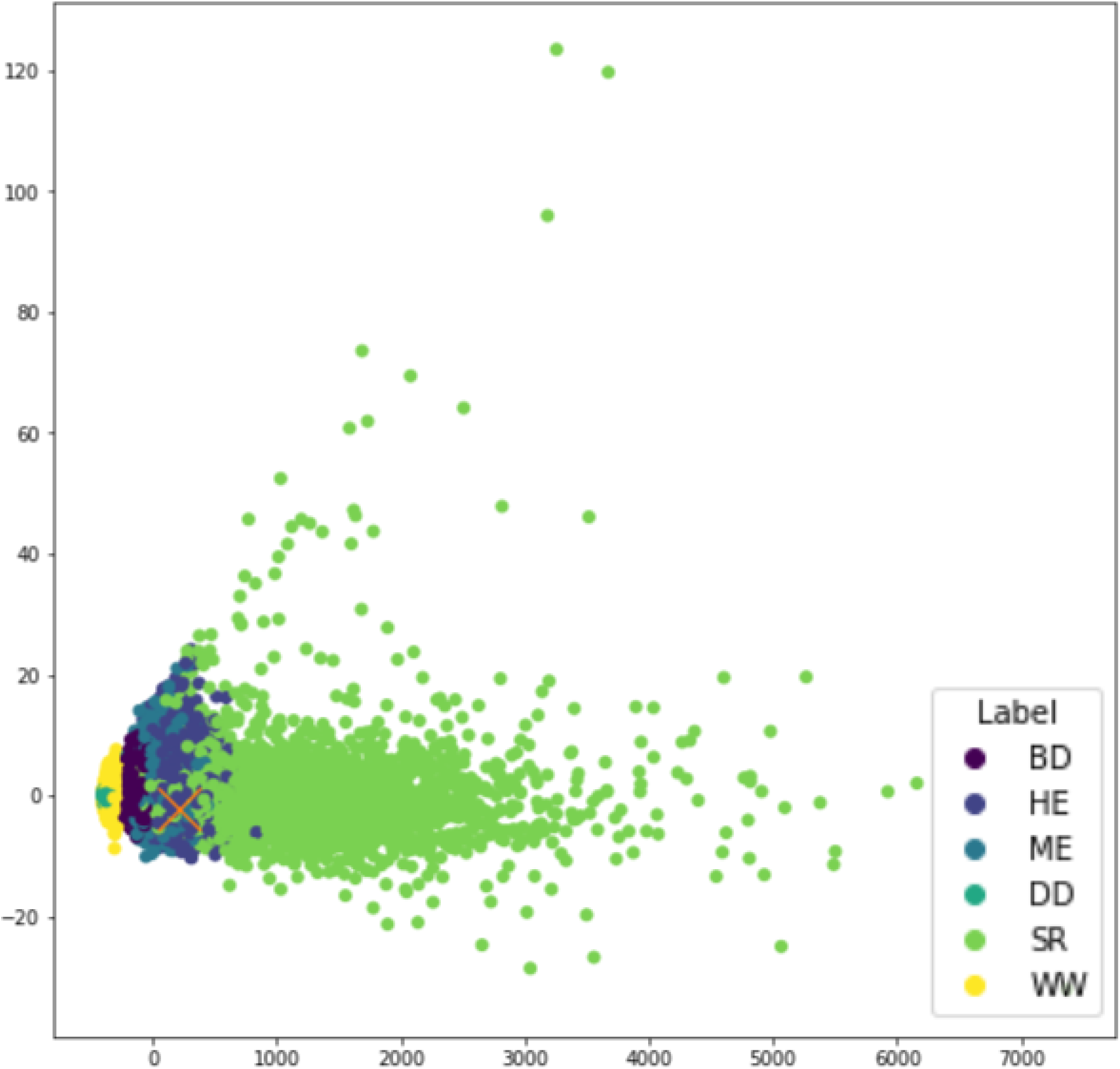
PCA performed over the vectorised trees (CDVm) of the 674 tips training dataset and its corresponding eucalyptus empirical case (represented as a red cross).

**Fig. 4.**
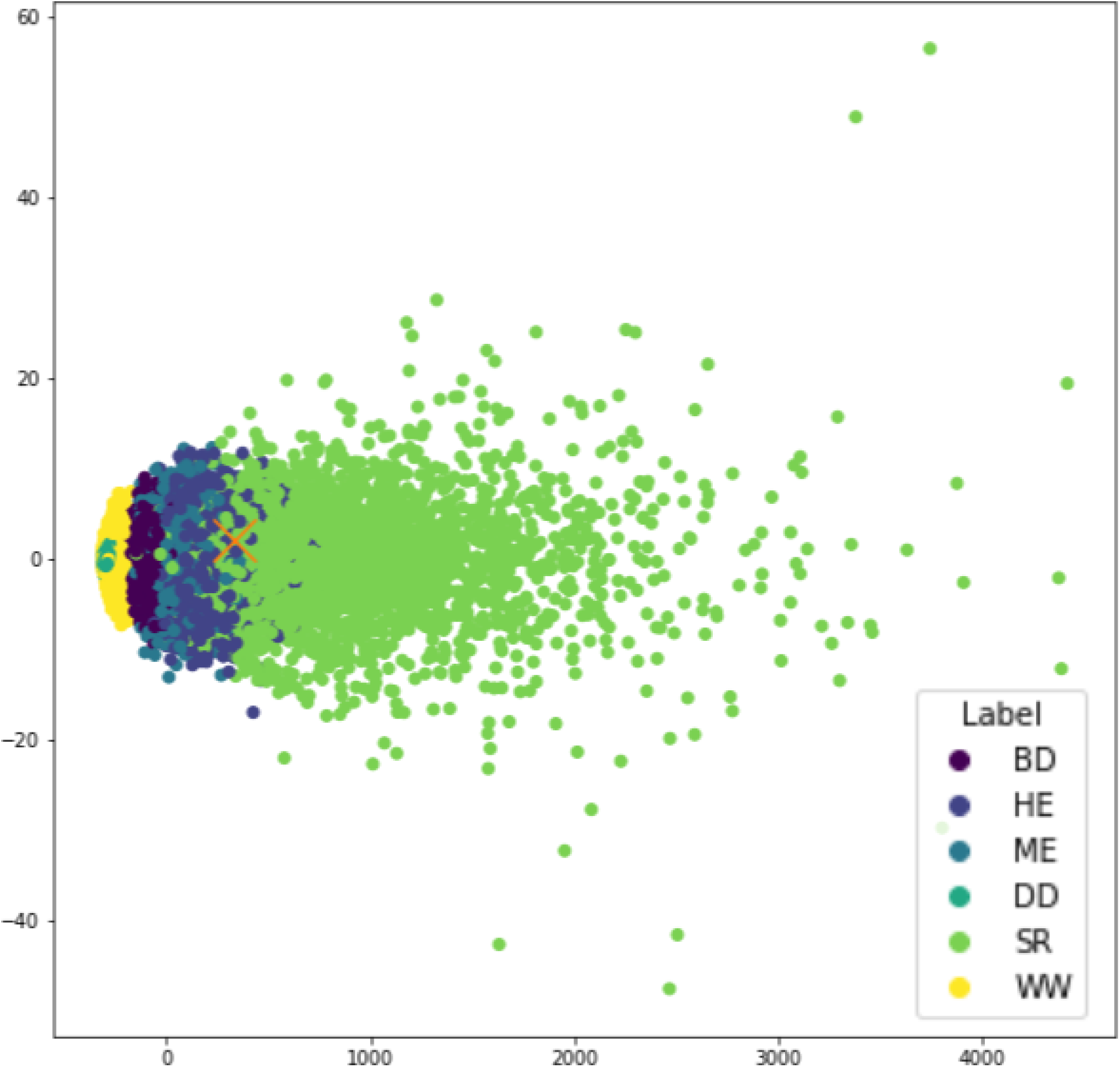
PCA performed over the vectorised trees (CDVm) of the 489 tips training dataset and its corresponding conifers empirical case (represented as a red cross).

**Fig. 5.**
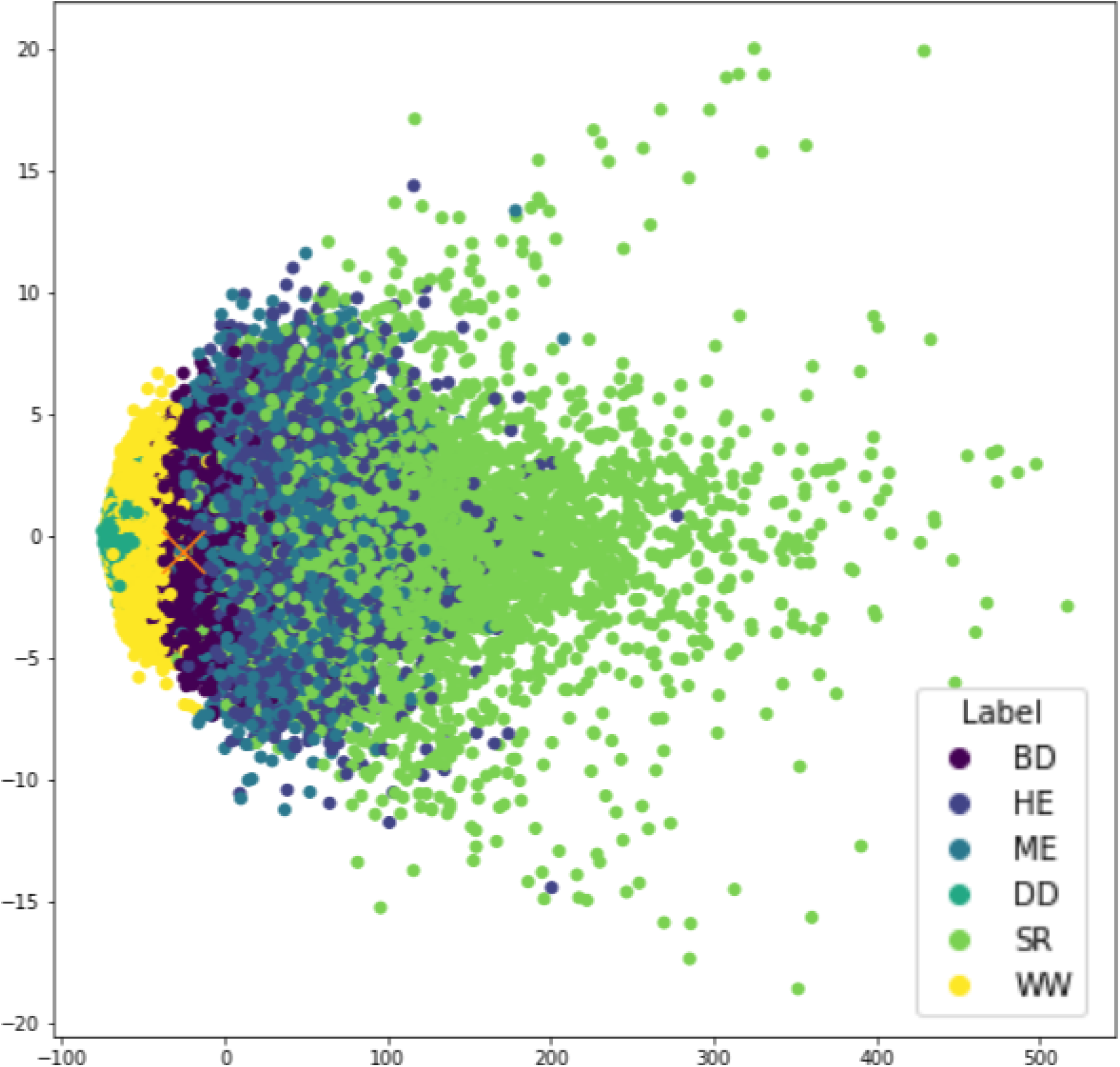
PCA performed over the vectorised trees (CDVm) of the 87 tips training dataset and its corresponding cetaceans empirical case (represented as a red cross).

**Fig. 6.**
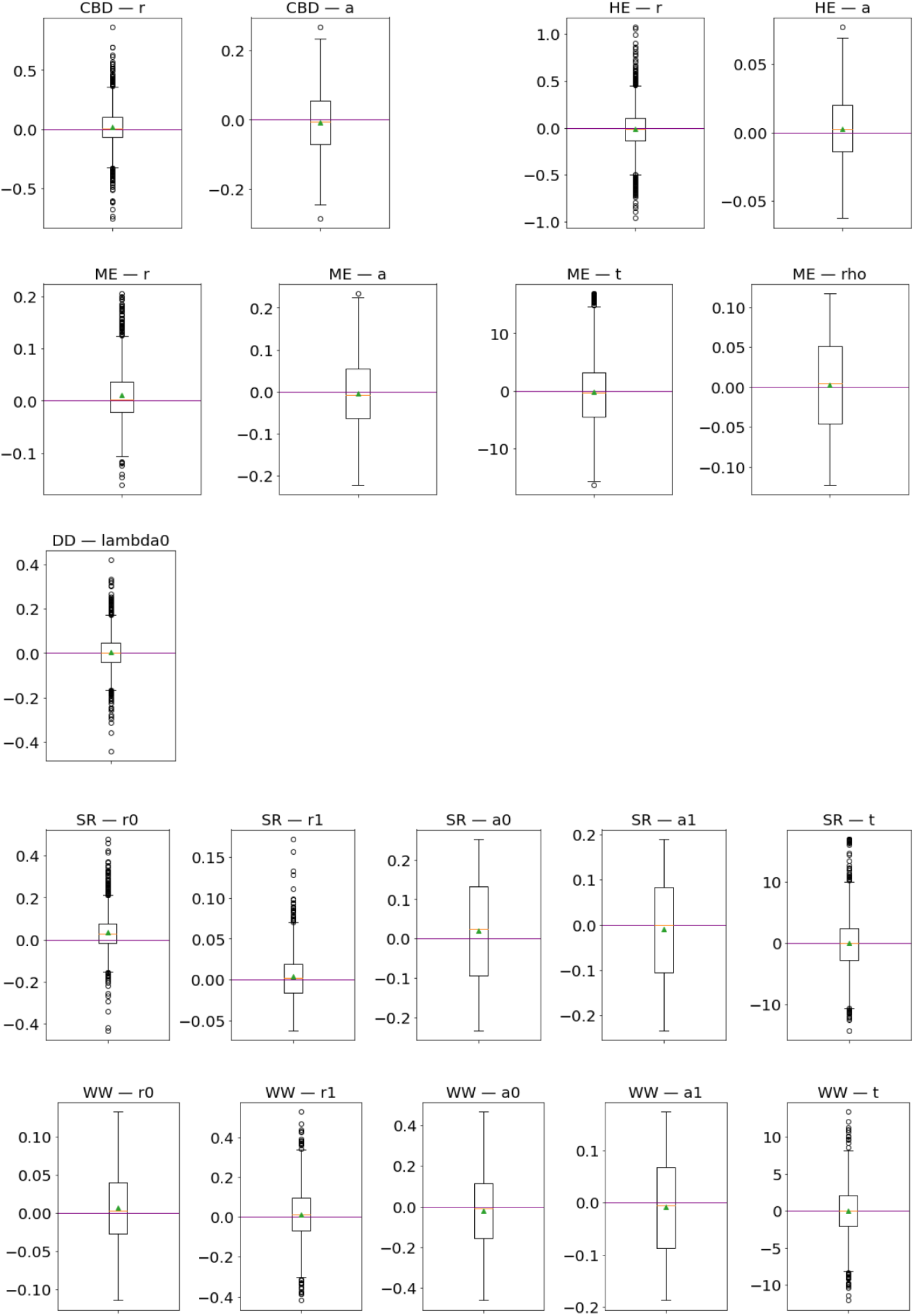
Box and whisker plots showing the prediction errors (*predicted_i_* − *real_i_*) across our set of DL regression models for the 674 tips dataset. The mean and the median are respectively represented as a green triangle and an orange line. The purple line marks the reference where predictions match target values exactly (error = 0). Definitions of the acronyms used for diversification scenarios and parameters can be found in Table 1 of the main text.

**Fig. 7.**
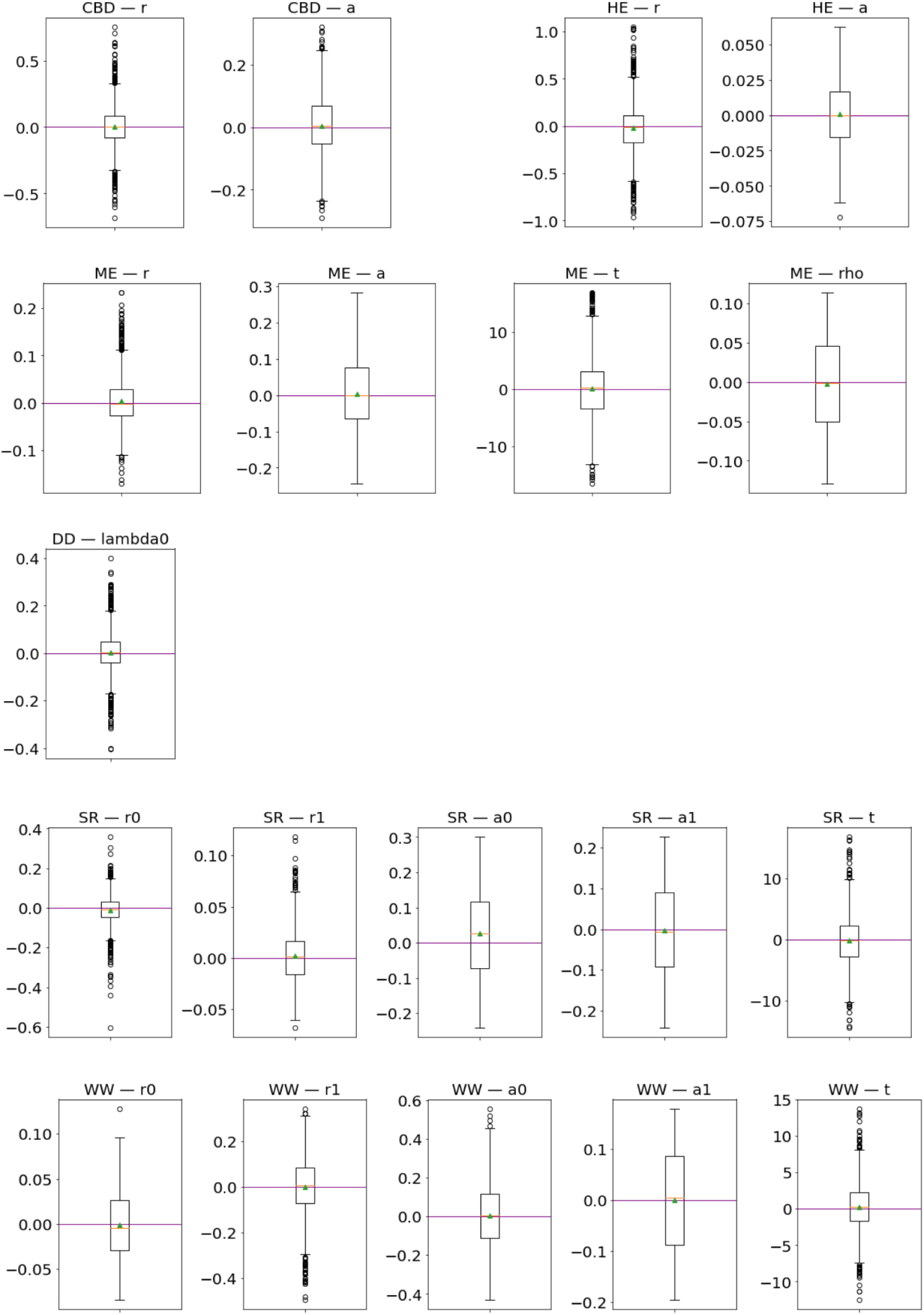
Box and whisker plots showing the prediction errors (*predicted_i_* − *real_i_*) across our set of DL regression models for the 489 tips dataset. The mean and the median are respectively represented as a green triangle and an orange line. The purple line marks the reference where predictions match target values exactly (error = 0). Definitions of the acronyms used for diversification scenarios and parameters can be found in Table 1 of the main text.

**Fig. 8.**
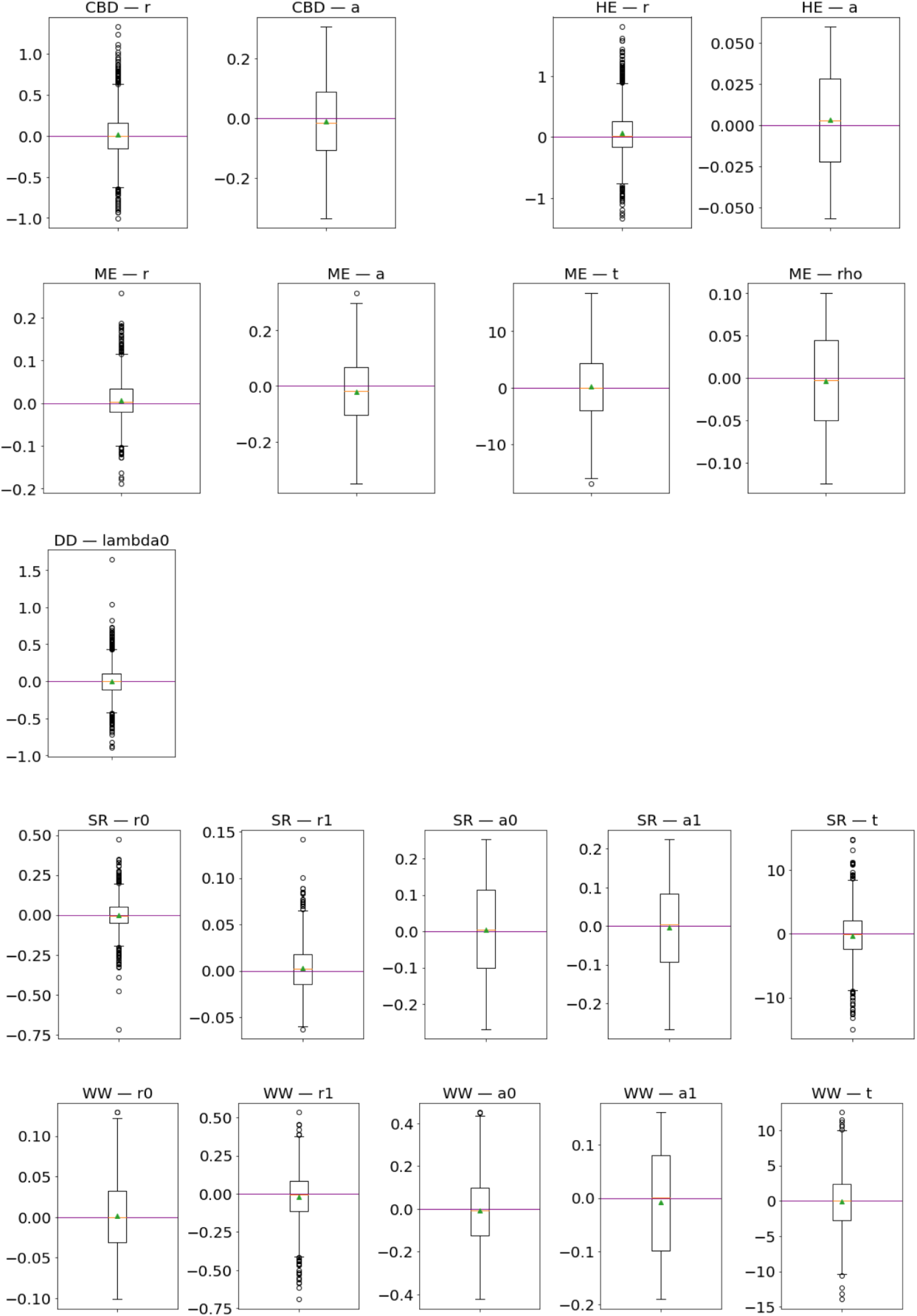
Box and whisker plots showing the prediction errors (*predicted_i_* − *real_i_*) across our set of DL regression models for the 87 tips dataset. The mean and the median are respectively represented as a green triangle and an orange line. The purple line marks the reference where predictions match target values exactly (error = 0). Definitions of the acronyms used for diversification scenarios and parameters can be found in Table 1 of the main text.

**Fig. 9.**
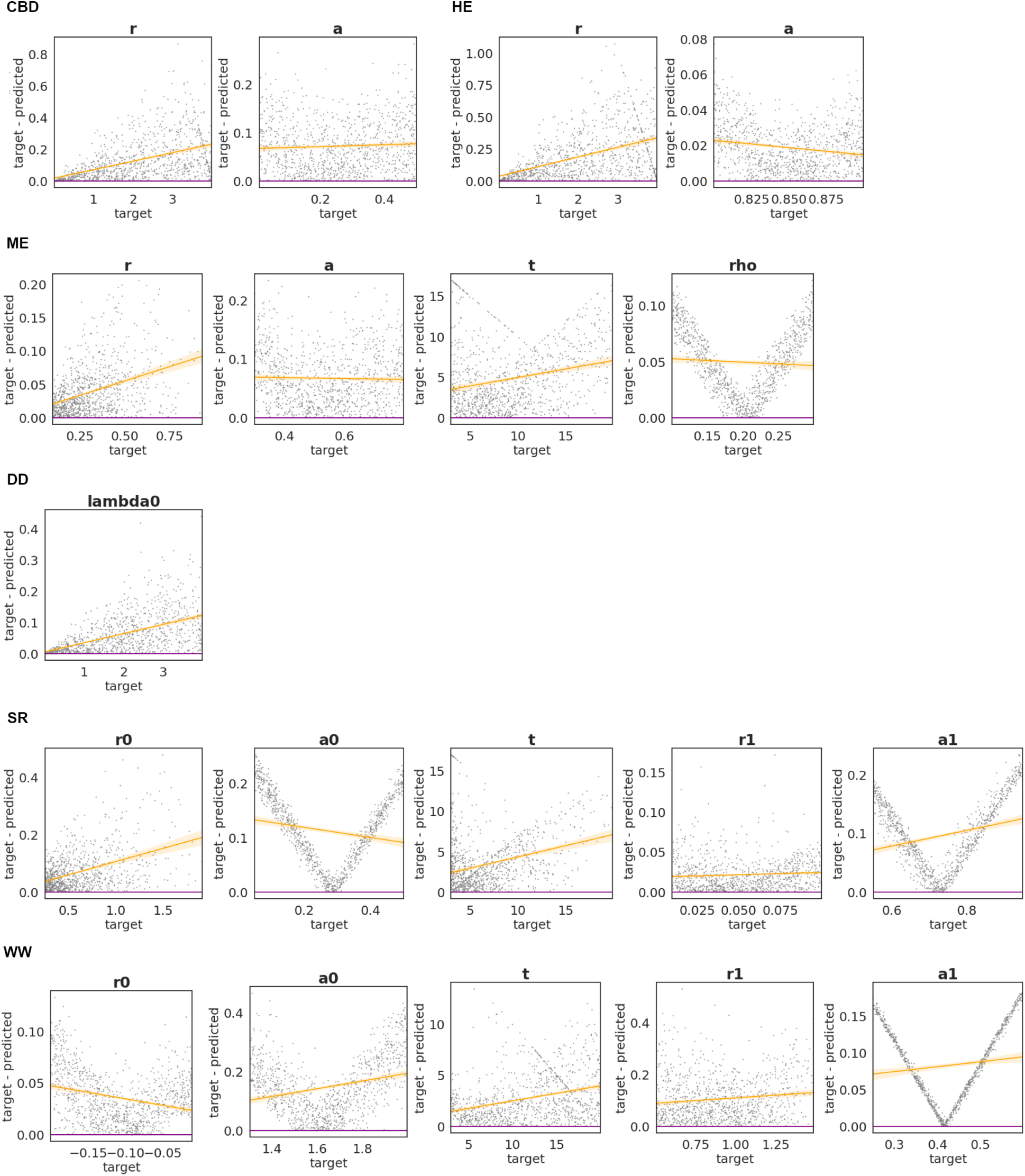
Absolute error versus target parameter values for DL regression models applied to each diversification scenario, based on 1,000 phylogenetic trees from the 674-tip simulated dataset. Each dot represents the error of a single prediction. The purple line marks the reference where predictions match target values exactly (error = 0), and the orange line shows the local polynomial regression with its 95% confidence interval.

**Fig. 10.**
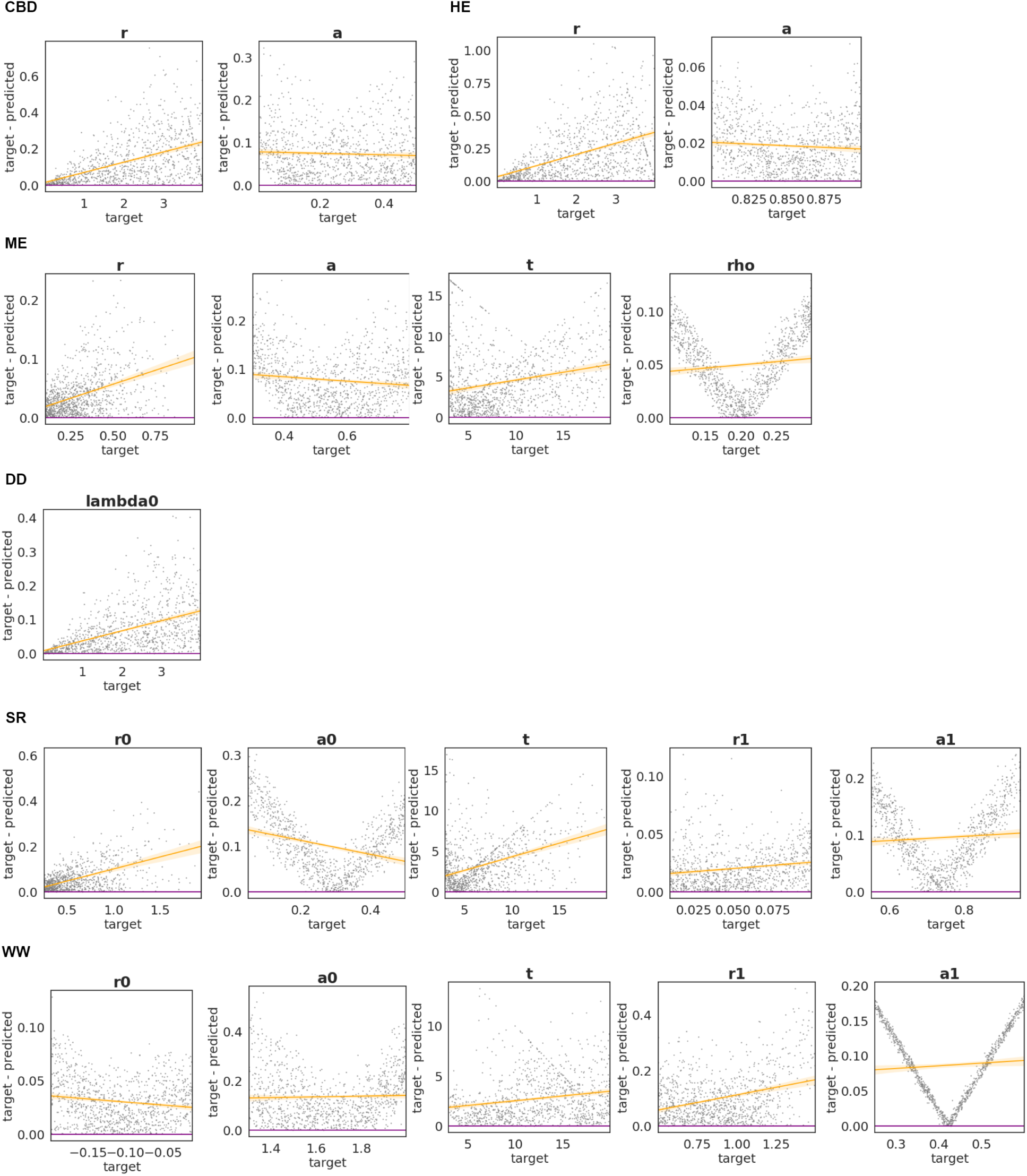
Absolute error versus target parameter values for DL regression models applied to each diversification scenario, based on 1,000 phylogenetic trees from the 489-tip simulated dataset. Each dot represents the error of a single prediction. The purple line marks the reference where predictions match target values exactly (error = 0), and the orange line shows the local polynomial regression with its 95% confidence interval.

**Fig. 11.**
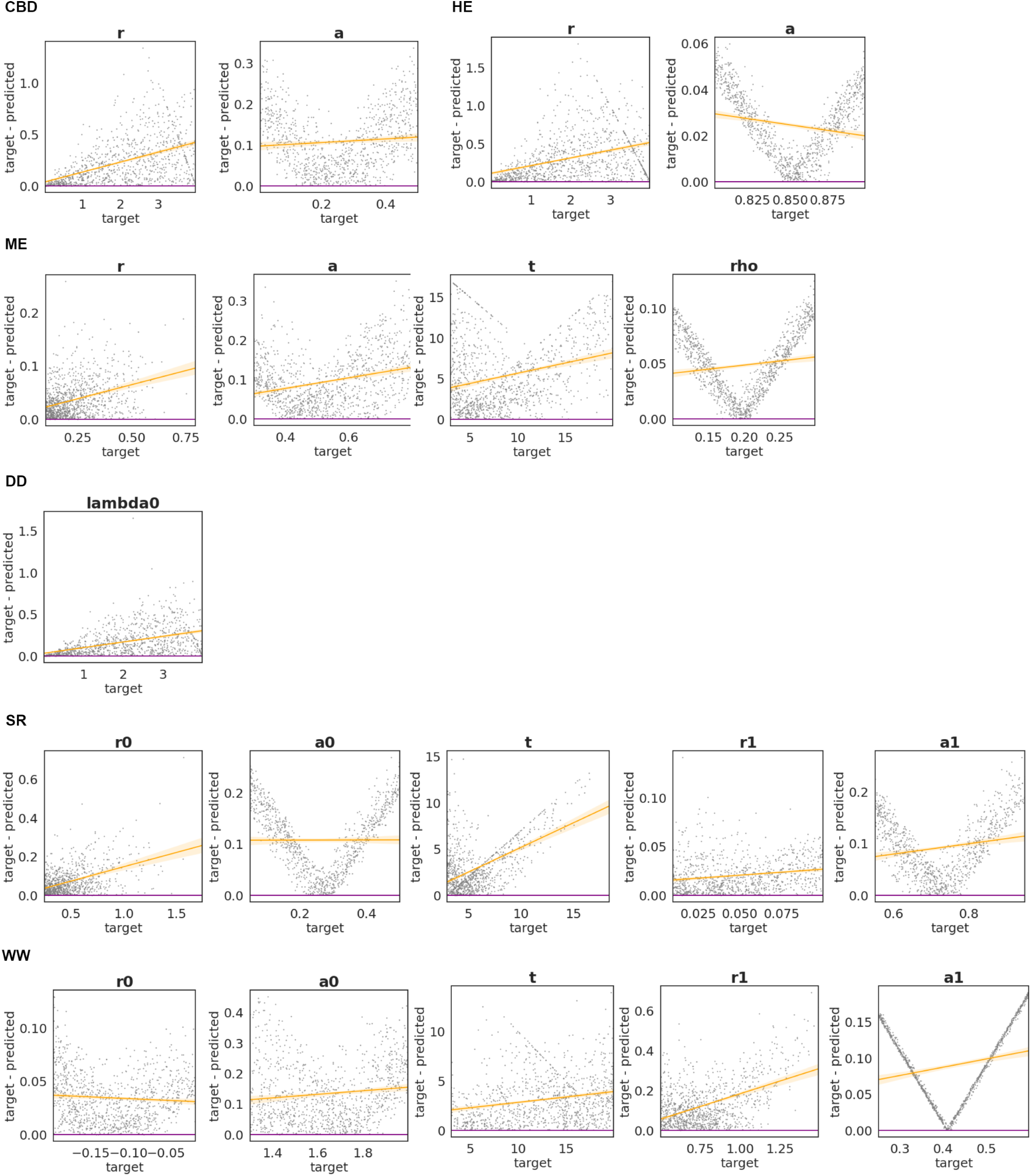
Absolute error versus target parameter values for DL regression models applied to each diversification scenario, based on 1,000 phylogenetic trees from the 87-tip simulated dataset. Each dot represents the error of a single prediction. The purple line marks the reference where predictions match target values exactly (error = 0), and the orange line shows the local polynomial regression with its 95% confidence interval.

**Fig. 12.**
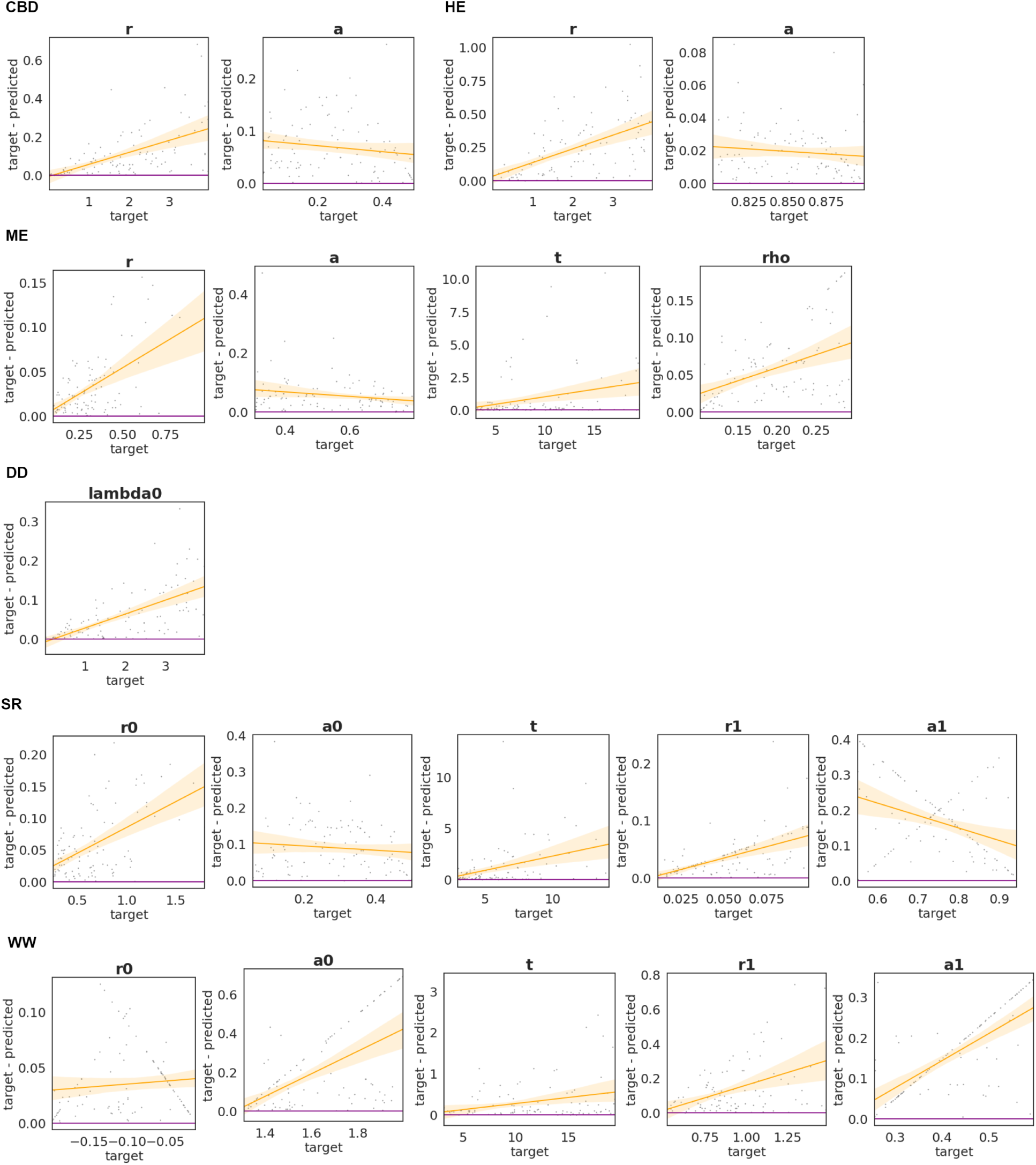
Absolute error versus target parameter values for MLE in each diversification scenario, based on 100 phylogenetic trees from the 674-tip simulated dataset. Each dot represents the error of a single prediction. The purple line marks the reference where predictions match target values exactly (error = 0), and the orange line shows the local polynomial regression with its 95% confidence interval.

**Fig. 13.**
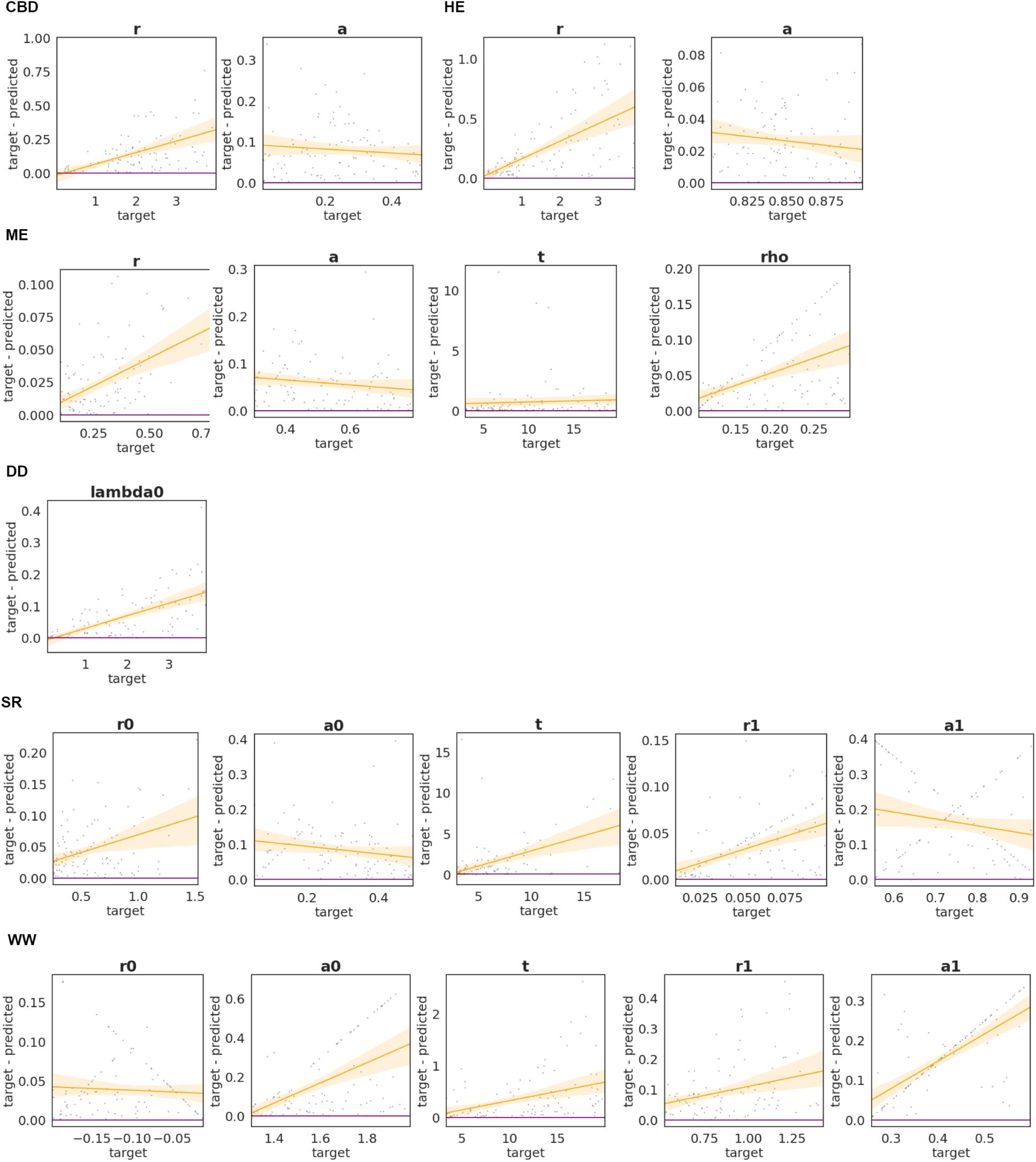
Absolute error versus target parameter values for MLE in each diversification scenario, based on 100 phylogenetic trees from the 489-tip simulated dataset. Each dot represents the error of a single prediction. The purple line marks the reference where predictions match target values exactly (error = 0), and the orange line shows the local polynomial regression with its 95% confidence interval.

**Fig. 14.**
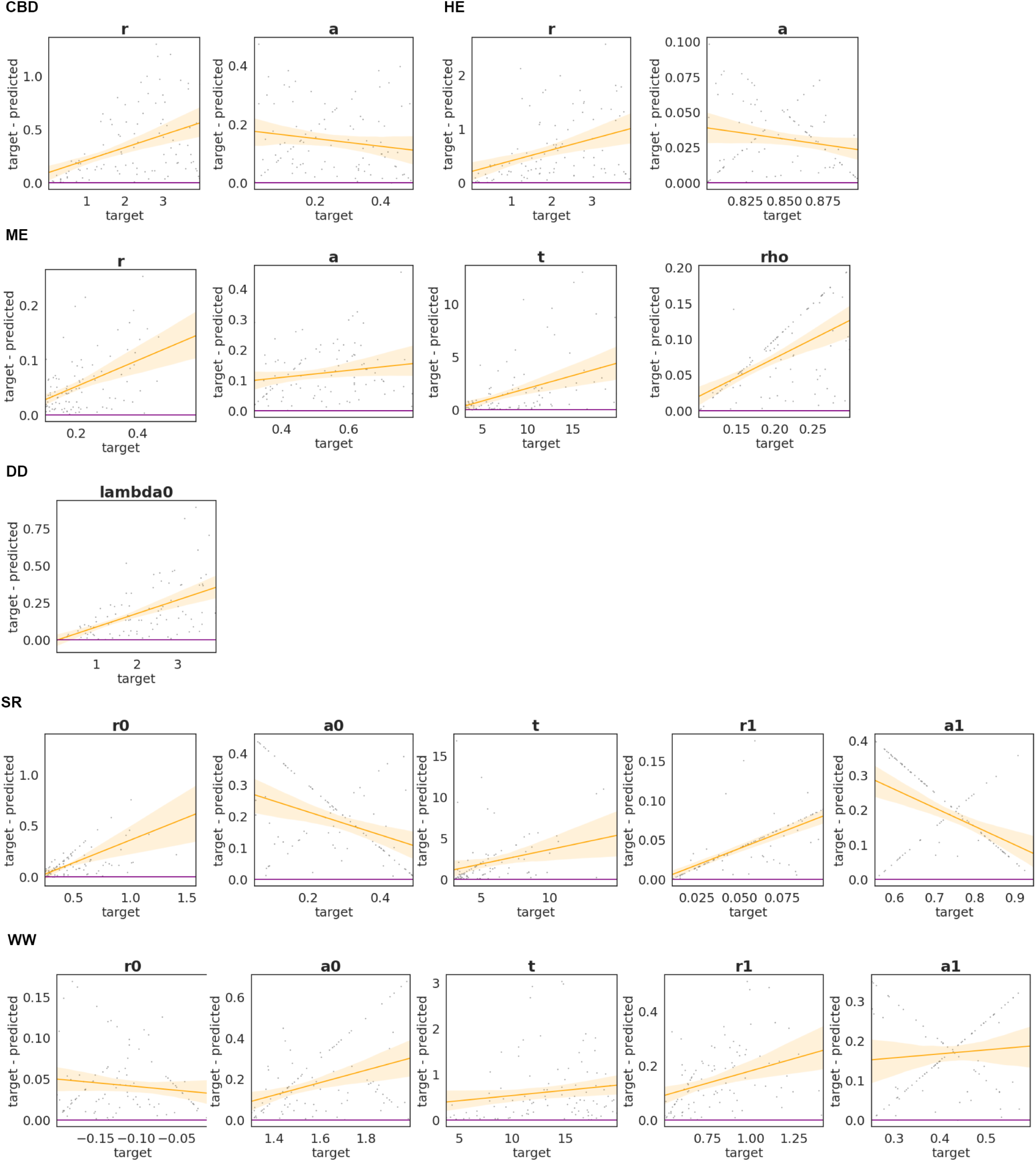
Absolute error versus target parameter values for MLE in each diversification scenario, based on 100 phylogenetic trees from the 87-tip simulated dataset. Each dot represents the error of a single prediction. The purple line marks the reference where predictions match target values exactly (error = 0), and the orange line shows the local polynomial regression with its 95% confidence interval.

**Table 7.** Classification results (accuracy, precision, recall, and F1-score) for our DL model trained on all simulation datasets combined (674 tips, 489 tips, and 87 tips).

|  | Accuracy | Precision | Recall | F1-Score |
| --- | --- | --- | --- | --- |
| All tips model | 0.8858 | 0.8865 | 0.8857 | 0.8855 |

**Table 8.** MAE values for our six regression models (one per diversification scenario), each trained on a combined dataset of simulated trees with 674, 489, and 87 tips.

| Parameter |  |  |  |  |  |  |
| --- | --- | --- | --- | --- | --- | --- |
| r0 | a0 | t | r1 | a1 | $\rho$ | $\lambda_0$ |
| CBD |  |  |  |  |  |  |
| 0.1685 | 0.0854 | - | - | - | - | - |
| HE |  |  |  |  |  |  |
| 0.2762 | 0.0208 | - | - | - | - | - |
| ME |  |  |  |  |  |  |
| 0.0358 | 0.0797 | 5.3608 | - | - | 0.0493 | - |
| DD |  |  |  |  |  |  |
| - | - | - | - | - | - | 0.1087 |
| SR |  |  |  |  |  |  |
| 0.0712 | 0.1088 | 3.0887 | 0.0208 | 0.0927 | - | - |
| WW |  |  |  |  |  |  |
| 0.0370 | 0.1464 | 2.5655 | 0.1169 | 0.0879 | - | - |

**Fig. 15.**
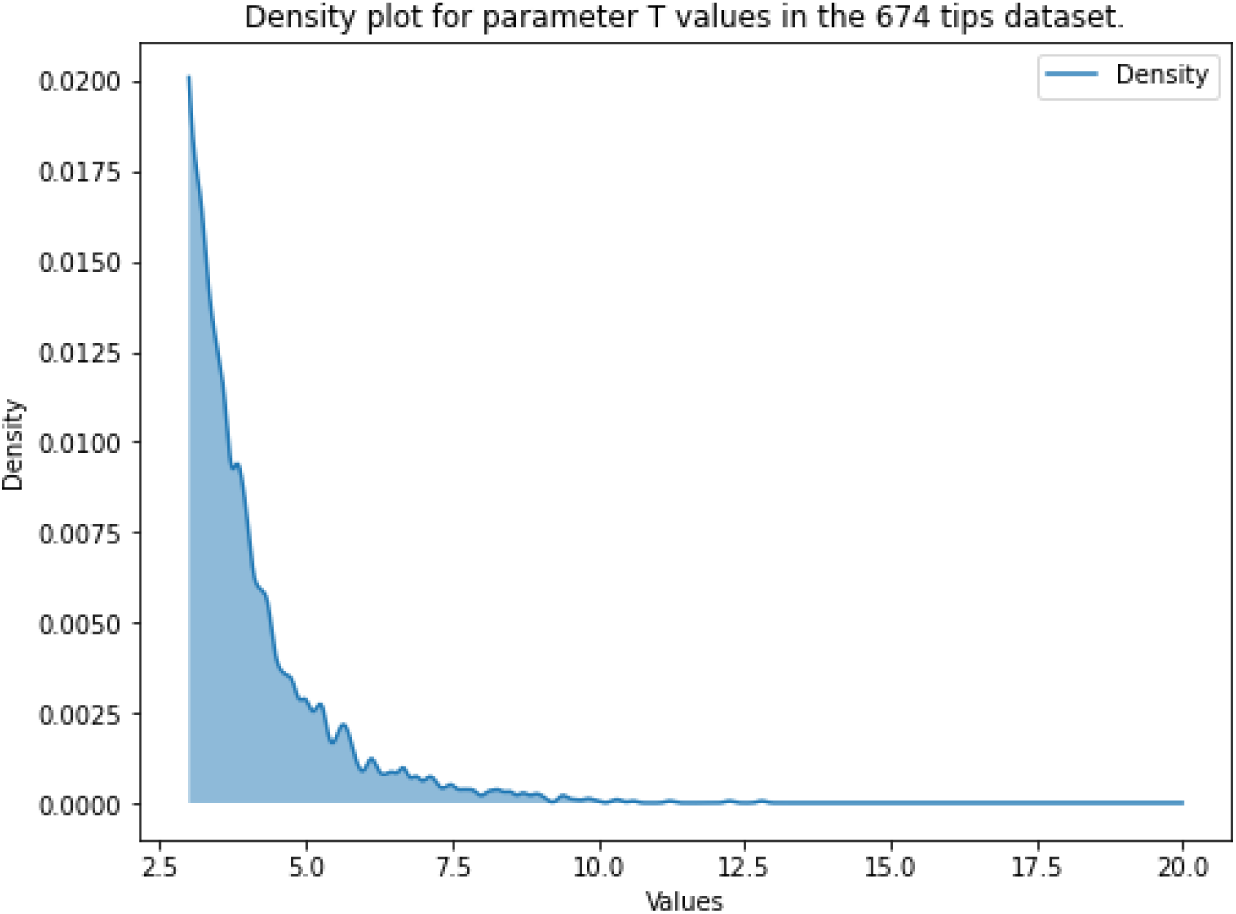
Density of parameter T values for the SR diversification scenario in the 674-tip simulation dataset. The density is calculated using Gaussian Kernel Density Estimation (KDE) and represented by a blue line.

**Fig. 16.**
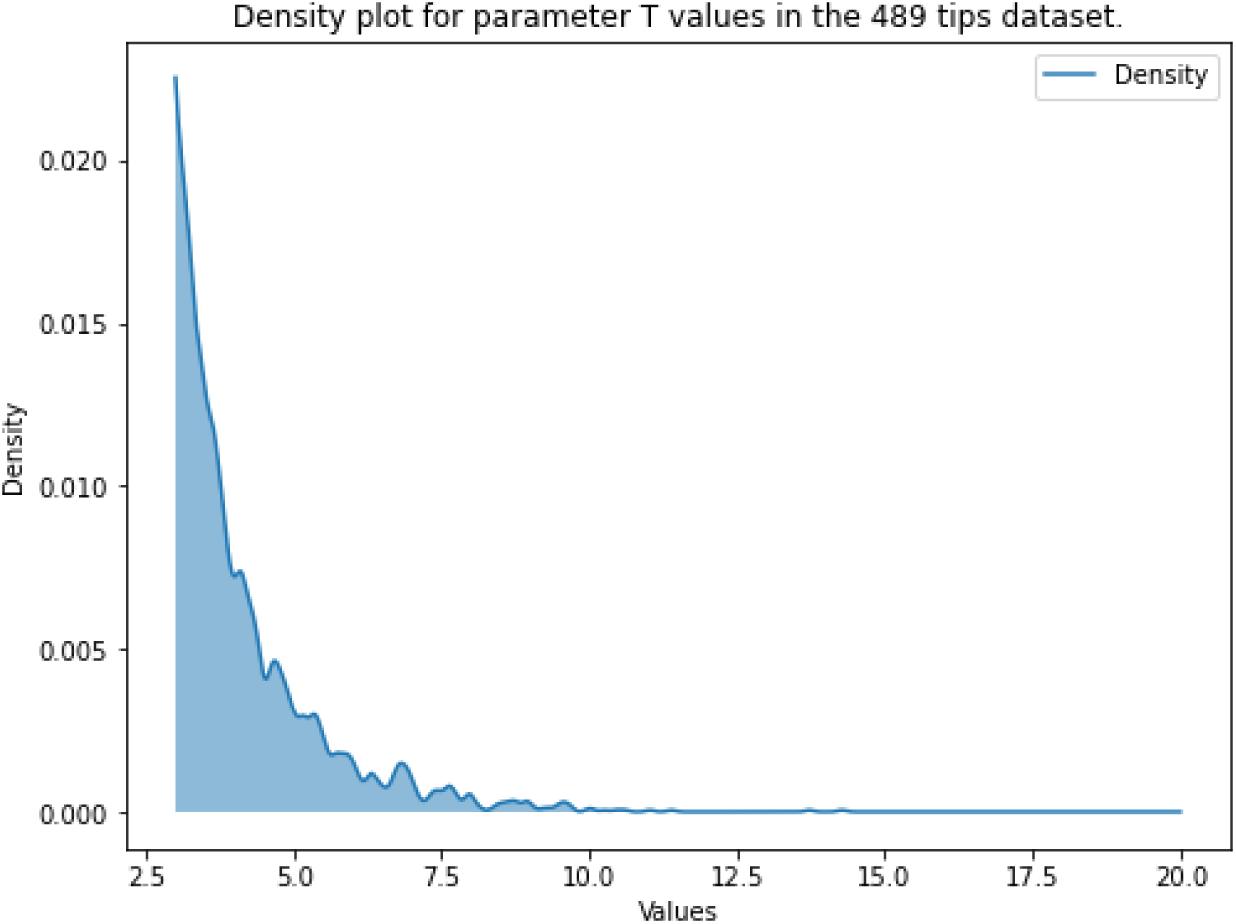
Density of parameter T values for the SR diversification scenario in the 489-tip simulation dataset. The density is calculated using Gaussian Kernel Density Estimation (KDE) and represented by a blue line.

**Fig. 17.**
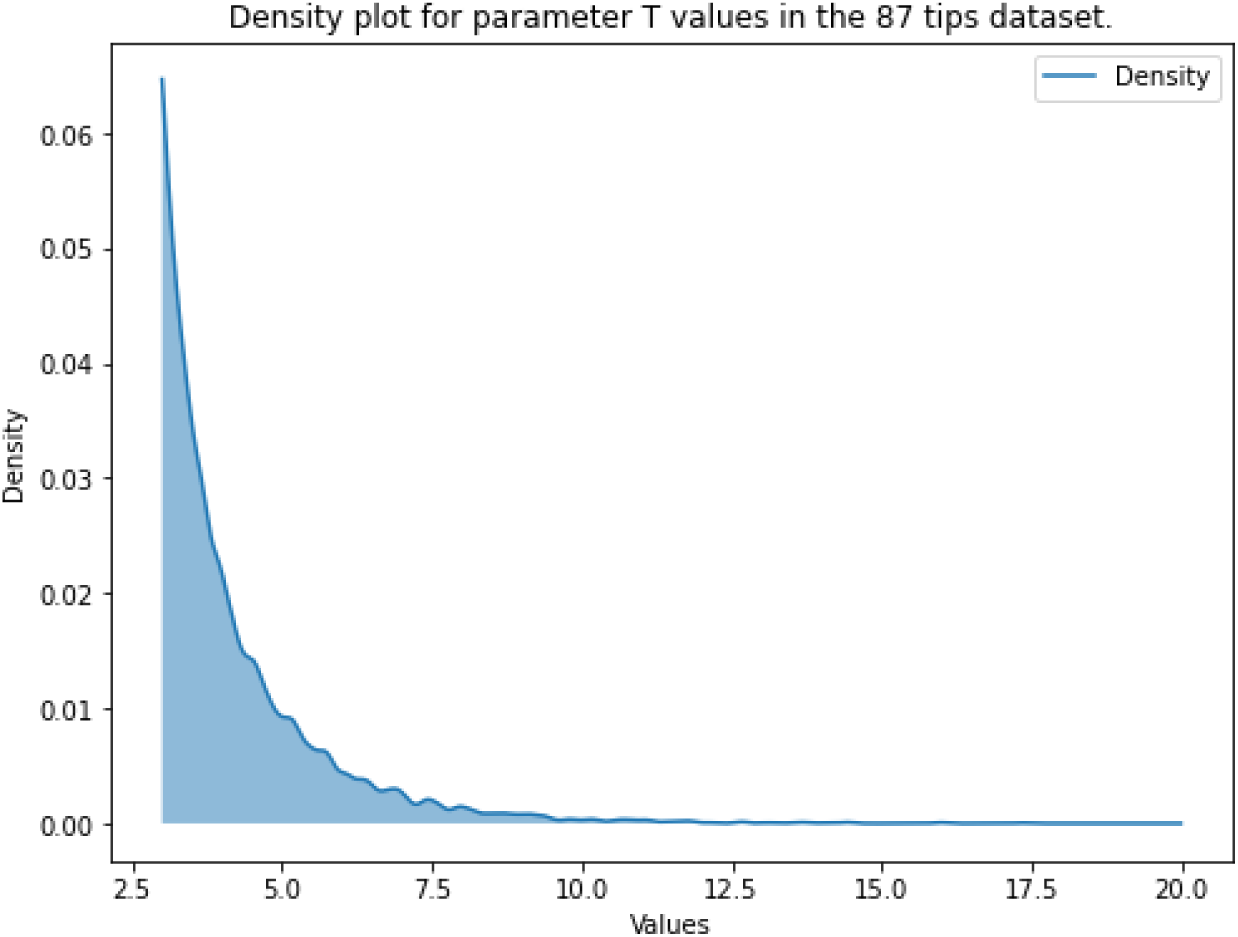
Density of parameter T values for the SR diversification scenario in the 87-tip simulation dataset. The density is calculated using Gaussian Kernel Density Estimation (KDE) and represented by a blue line.

**Fig. 18.**
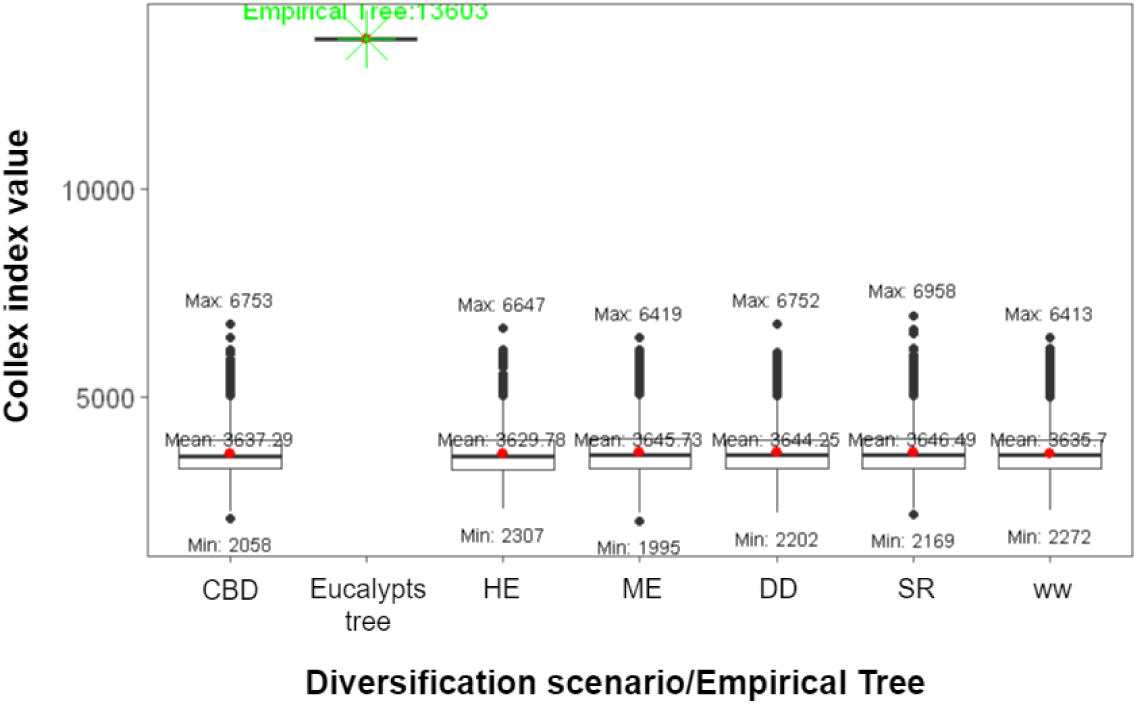
Boxplot representing the colless index values for the phylogenetic trees simulated under the different diversification scenarios in the 674 tips dataset. The green cross represents the value of the colless index for the empirical phylogeny of eucalypts.

**Fig. 19.**
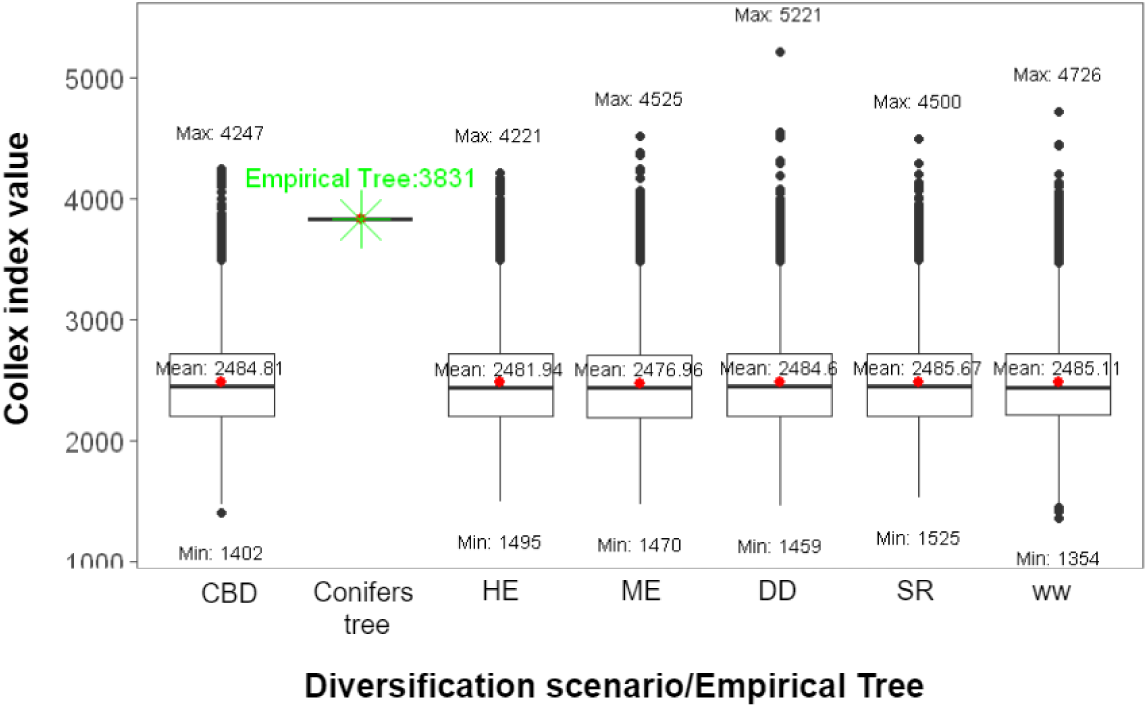
Boxplot representing the colless index values for the phylogenetic trees simulated under the different diversification scenarios in the 489 tips dataset. The green cross represents the value of the colless index for the empirical phylogeny of conifers.

**Fig. 20.**
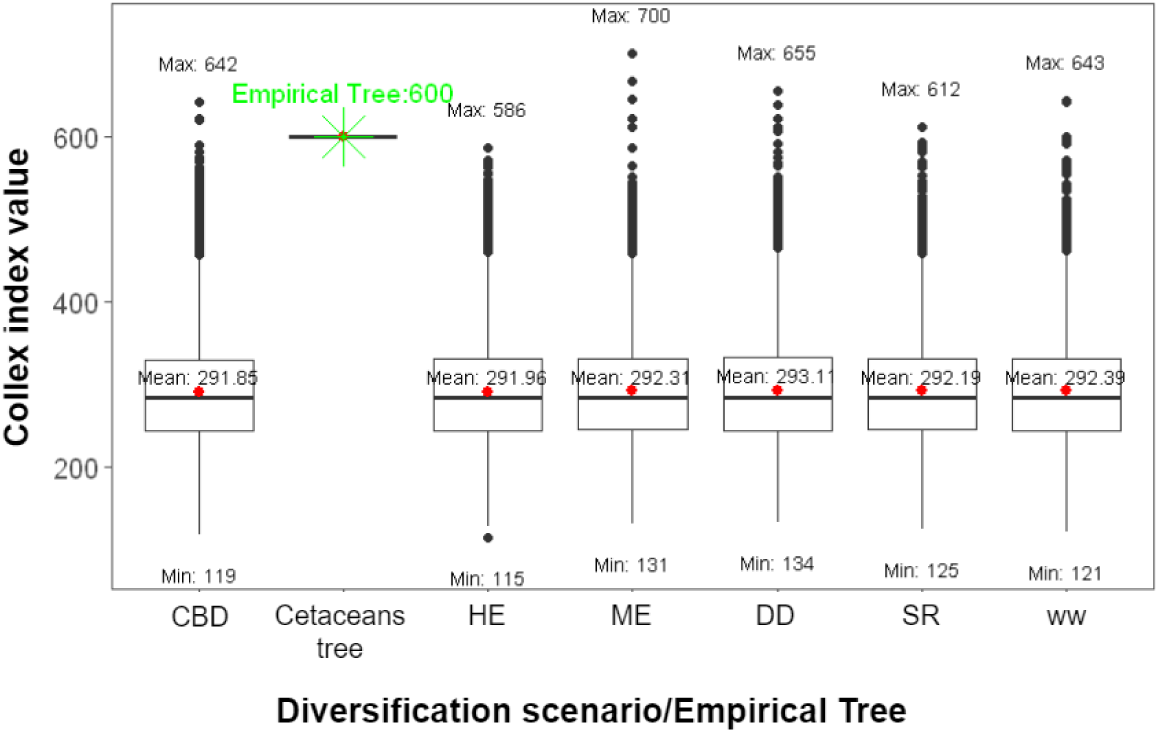
Boxplot representing the colless index values for the phylogenetic trees simulated under the different diversification scenarios in the 87 tips dataset. The green cross represents the value of the colless index for the empirical phylogeny of cetaceans.

## Notes

https://github.com/pablogpena/deep_birth_death

### Competing Interest Statement

The authors have declared no competing interest.

### Summary of Updates

Author name corrected, all of the rest is the same until next revisions.

